# Direct and ultrasensitive bioluminescent detection of intact respiratory viruses

**DOI:** 10.1101/2024.09.22.614338

**Authors:** Alexander Gräwe, Harm van der Veer, Seino A.K. Jongkees, Jacky Flipse, Iebe Rossey, Robert P. de Vries, Xavier Saelens, Maarten Merkx

## Abstract

Respiratory viruses such as SARS-CoV-2, influenza, and respiratory syncytial virus (RSV) represent pressing health risks. Rapid diagnostic tests for these viruses detect single antigens or nucleic acids, which do not necessarily correlate with the amount of intact virus. Instead, specific detection of intact respiratory virus particles may better assess the contagiousness of a patient. Here, we report GLOVID, a modular biosensor platform to detect intact virions against a background of ‘free’ viral proteins in solution. Our approach harnesses the multivalent display of distinct proteins on the surface of a viral particle to template the reconstitution of a split luciferase, allowing specific, single-step detection of intact influenza A and RSV virions corresponding to 0.1 - 0.3 fM of genomic units. The protein ligation system used to assemble GLOVID sensors is compatible with a broad range of binding domains, including nanobodies, scFv fragments and cyclic peptides, which allows straightforward adjustment of the sensor platform to target different viruses.

## MAIN TEXT

Respiratory viruses represent a continuous challenge to healthcare systems and a potential source of new pandemics ^1,2^. Influenza A virus (IAV) is under constant evolutionary pressure to escape recognition by the human immune system and has a long history of pandemics in healthy populations, whereas RSV is another clinically relevant respiratory virus causing severe illness in infants and the elderly. Testing individuals at the front line is essential during a virus outbreak as it can prevent uncontrolled virus spread and avoid healthcare system overload. Current point-of-need tests measure nucleic acids or protein antigens ^3,4^. However, these virus components do not necessarily correlate with the amount of infectious virus particles in a sample ^5,6^. Therefore, we aimed to develop a sensor platform that allows sensitive and specific detection of intact virus particles, where the challenge is to translate the presence of intact virus particles into a sensitive optical signal that can be readily detected in the sample of interest ^6,7^.

The GLOVID (GLOwing Virus Detection) sensor platform presented here harnesses the multivalent display of distinct proteins on the surface of a viral particle to template the reconstitution of a split luciferase. Homogeneous bioluminescent assays based on proximity-driven complementation of a split luciferase have several advantages for point-of-need assays, including an intrinsic high dynamic range (100-to 1000-fold increase in signal) and straightforward sensor design ^8,9^. Since bioluminescence does not require excitation, such assays typically have very low background and can be conducted directly in complex samples, using a simple digital camera or smartphone for detection ^10,11^. A recent example of a proximity-driven split luciferase sensor platform is RAPPID, a homogeneous sandwich immunoassay format for detecting protein biomarkers that uses the split luciferase NanoBiT ^12^. In NanoBiT, complementation of a large (LgBiT) and a small (SmBiT) subunit is required to form an active luciferase complex ^13^.

A prerequisite for GLOVID is modularity, so that binding domains can be easily exchanged to adapt to the rapid evolution of new viral strains. Here, we show that the recently described DogTag/DogCatcher protein ligation system ^14^ enables straightforward construction of NanoBiT-based sensor components. The DogTag peptide forms a spontaneous isopeptide bond with DogCatcher upon mixing, allowing assembly of DogTagged binders and split sensors with (multiple) DogCatcher domains into branched protein structures. Since many viral surface proteins are present as multimers, the GLOVID platform was designed to enable the functionalization of the LgBiT and SmBiT components with either one or three binding domains. Post-translational binder conjugation using the DogTag/DogCatcher allows efficient synthesis of multivalent binders and is shown to be compatible with a broad range of binders, including nanobodies, single-chain variable fragments (scFv) domains and synthetic cyclic peptides.

The modularity of the GLOVID platform enables a broad range of assay formats, ranging from the detection of single proteins and protein trimers to the specific detection of intact viruses by targeting two different proteins on the viral surface (**Figure 1A**). The latter format allows specific detection of intact virus particles only.

**Figure 1.**
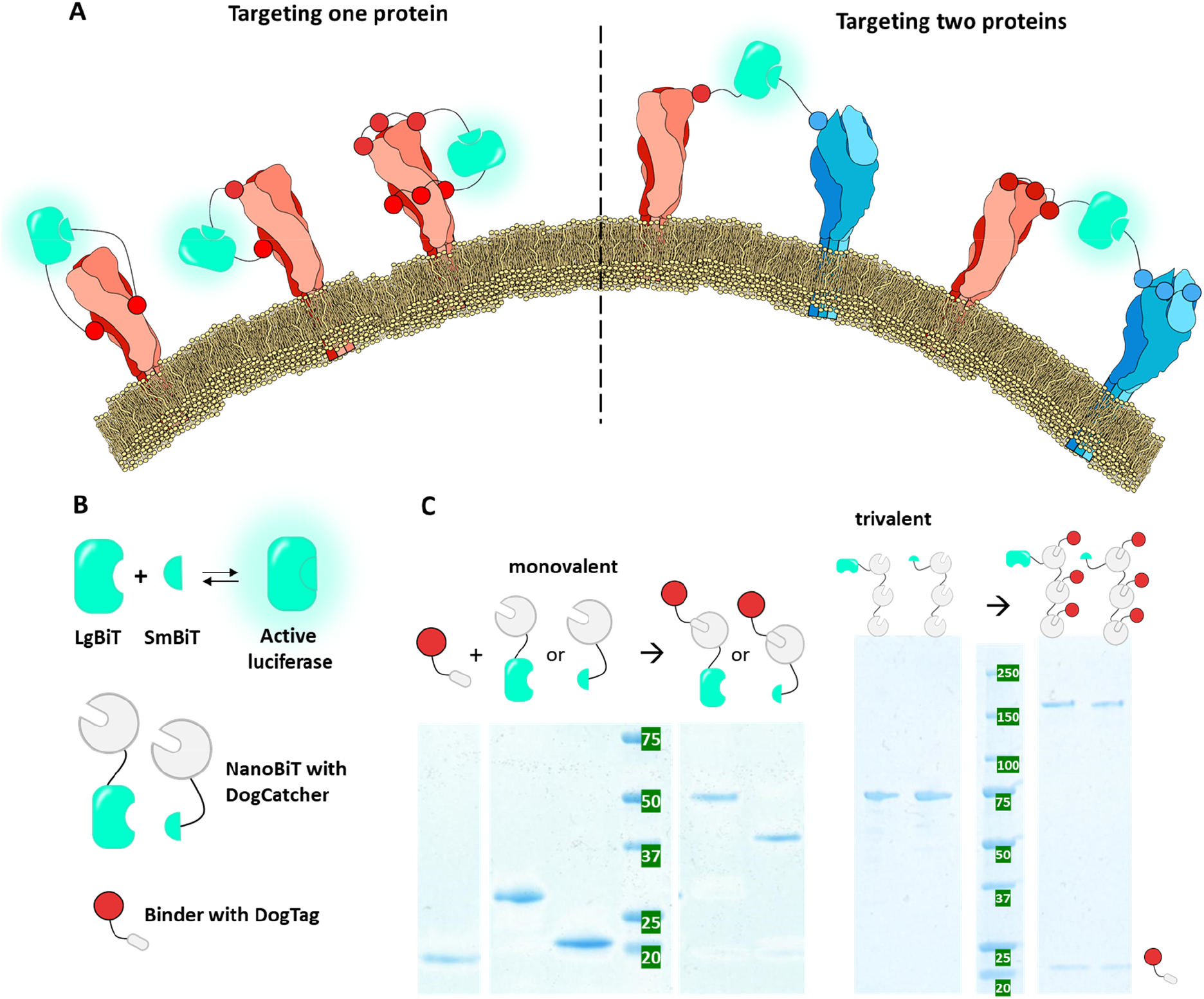
Concept and assembly of the GLOVID platform. A) Sensor platform designs. The split luciferase NanoBiT is fused to specific binders that target viral surface proteins via mono- or multivalent interactions. Complementation of the split system is required to generate a signal. Conceptually, targeting two different surface proteins makes the output signal dependent on intact virions, while targeting one surface protein may also lead to the detection of soluble proteins. B) Toolbox to combine NanoBiT with antigen-specific binders. The NanoBiT components LgBiT and SmBiT that, upon complementation, form an active luciferase are fused to DogCatcher (grey circle), whereas target-specific binders are fused to DogTag (grey oval). C) SDS-PAGEs showing successful assembly of monovalent and trivalent sensor components, exemplified for DogTagged nanobody SD36 (red). Upon mixing, DogTag spontaneously forms an isopeptide bond with DogCatcher present in LgBiT-Dog1, SmBiT-Dog1, LgBiT-Dog3 and SmBiT-Dog3, leading to stably connected fusion proteins with a higher molecular weight. Monovalent systems were mixed in an equimolar ratio, whereas trivalent systems were mixed in a 4:1 DogTag:DogCatcher ratio to achieve complete conversion of DogCatcher.

## Results

### Design of the GLOVID platform

We aimed to create a versatile bioluminescent immunodiagnostics platform that allows attachment of various binders, including nanobodies, scFvs, and synthetic peptides. The affinity of these binders to their targets is a crucial determinant of the platform’s sensitivity. Since many viral proteins are present as homotrimers on the viral surface, the possibility of incorporating three binding domains was included to enhance avidity (**Figure 1A**). However, recombinant expression of fusion proteins with multiple copies of a binding domain is not trivial for disulfide-containing binding domains such as nanobodies and scFvs. In addition, genetic fusions to N- and C-termini of the binding domains, which is inevitable in genetic fusions, can affect their binding properties. We therefore chose a modular strategy in which binders are expressed and purified as individual domains, followed by posttranslational conjugation to LgBiT and SmBiT using the DogCatcher/DogTag system (**Figure 1A**). The platform is constructed by genetic fusion of LgBiT and SmBiT domains to one or three DogCatcher domains that can in turn be labeled with DogTagged binders (**Figure 1B**). We chose the SmBiT101 variant as its low binding affinity to LgBiT (*K*_*D*_ = 2.5 µM) should minimize background signal at low nM sensor concentrations. We selected the DogTag/DogCatcher system over the SpyTag/SpyCatcher system as the former works in a broader range of protein ligation scenarios ^14^, e.g., in cases where the DogTag needs to be inserted in a loop region of a binder because one or both termini of the binder are very close to the binding site ^15^. The DogCatcher domains were connected by 42 amino acid long, threonine-proline linkers (TP)_21_. These linkers are inspired by natural linkers ^16,17^ and were selected because they are flexible enough to adapt to different multivalent protein architectures, while their semi-flexible nature also allows them to effectively bridge distances between neighboring epitopes ^18,19^.

### GLOVID allows sensitive detection of soluble IAV hemagglutinin

As a first application to test the performance of the GLOVID platform, we chose the IAV hemagglutinin (HA)-trimer protein as a target. This HA-trimer is abundantly present on the IAV surface but can also be expressed as a soluble trimeric protein by fusion of the HA sequence to a GCN4 trimerization domain ^20^. Several well-characterized HA-binders have been reported, two of which were chosen here: SD36 and HSB2.A. SD36 is a nanobody that targets a conserved epitope in the HA stem region of IAV group 2 strains and binds A/Hongkong/1/1968/H3N2 HA (H3HK) with a high monovalent affinity (*K*_*D*_ = 2.4 nM) ^21^. HSB2.A is a small and stable *de novo* designed binding domain that targets the receptor binding site on the head region of IAV H3HK ^22^. Notably, it has been reported that the monovalent affinity of HSB2 is relatively weak (*K*_*D*_ > 50 nM) but can be substantially enhanced by trimerization ^22^.

Both the SD36 and HSB2.A fused with DogTags and the four GLOVID scaffold domains (LgBiT-Dog1, LgBiT-Dog3, SmBiT-Dog1, and SmBiT-Dog3) were obtained in good yields following recombinant expression in *E. coli* and purified using Ni-NTA and/or streptactin affinity chromatography. Next, GLOVID sensors were obtained by overnight incubation at 22 °C of the LgBiT-Dog1/SmBiT-Dog1 and LgBiT-Dog3/SmBiT-Dog3 with 1 and 4 equivalents of DogTagged binding domains, respectively. SDS-PAGE analysis confirmed complete functionalization of the monovalent and trivalent scaffold proteins (**Figure 1C**), demonstrating the efficient formation of the isopeptide bond between DogTag and DogCatcher. The ligation products were used in subsequent assays without further purification, as no unreacted scaffold domains were present.

First, we tested the performance of an assay in which SD36 was conjugated to both LgBiT-Dog1 and SmBiT-Dog1 (**Figure 2A**). Titration of trimeric H3HK protein to a mixture of 1 nM of each sensor component showed an 1100-fold increase of bioluminescent signal over a broad concentration range between 1 pM and 10 nM, corresponding to a limit of detection (LoD) of 3.7 pM (**Figure 2A**). Very low background bioluminescence was observed when no binders were attached to GLOVID components (Figure S1). A ‘hook’ effect was observed for higher target concentration, corresponding to a lower probability of simultaneous binding of SmBiT and LgBiT components to one trivalent target. The large increase in luminescence over a broad concentration range represents a substantial improvement compared to a recently reported single-component Dark-LUX bioluminescent sensor protein based on the same SD36-binding domains, which displayed a 5-fold increase in bioluminescent intensity and a LoD of 80 pM ^23^. The GLOVID assay using trivalent SD36-functionalized sensor components showed three orders of magnitude reduced sensitivity (LoD = 3.3 nM) compared to the monovalent version and a reduced dynamic range of bioluminescent signal (DR 49-fold) (**Figure 2B)**. This result is expected as binding of one of the trivalent sensor components to the three binding sites on an HA trimer would effectively block binding of the complementary sensor component to the same target. The observation that complementation is only observed at high concentrations of H3HK could be explained by the formation of complexes containing two HAs.

**Figure 2.**
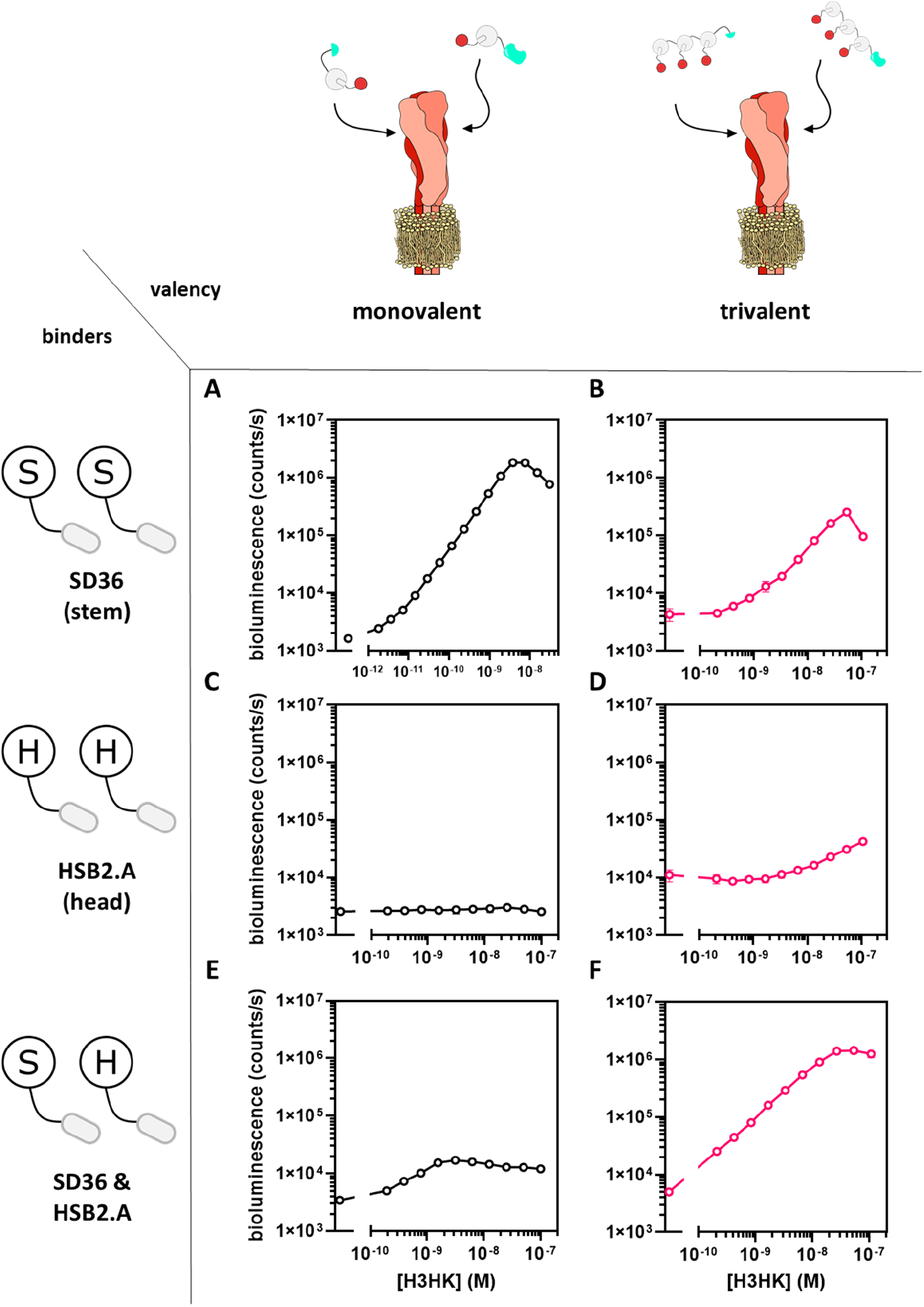
GLOVID titrations with H3HK hemagglutinin. A) Assay with SD36-functionalized, monovalent sensor components. B) Assay with SD36 functionalized, trivalent sensor components. C) Assay with HSB2.A functionalized, monovalent sensor components. D) Assay with HSB2.A functionalized, trivalent sensor components. E) Assays with monovalent combinations of SD36 (stem binder) and HSB2.A (head binder). F) Assay with a combination of trivalent SD36-SmBiT and trivalent HSB2.A-LgBiT. Experimental conditions: A) Final concentration of GLOVID parts 1 nM, 1xPBS + 1mg/ml BSA, final NanoGlo dilution of 1:2000, 1 h incubation at 22 °C. C) & E), as in A) but with 16 h incubation at 4 °C. B), D) & F): As in A) but with a final concentration of GLOVID parts of 2 nM for 16 h incubation at 4 °C. Error bars correspond to the standard deviation of n=3 replicates. Lines are used to connecting the points for better visualization.

GLOVID sensors containing the head-binding HSB2.A-domain showed no (monovalent system; **Figure 2C**) or only a very minor (trivalent; **Figure 2D**) increase in luminescent signal upon addition of H3HK. The non-responsiveness of the monovalent system can be explained by the low affinity of the monovalent HSB2.A:H3HK interaction, being too weak to support sensor complementation at the H3HK head region. This interaction is expected to be enhanced for the trivalent system, but since the complementary sensor components target the same trivalent binding site, complementation is still largely prevented here as well.

Finally, we tested a GLOVID setup in which the LgBiT and SmBiT domains were conjugated to different binding domains. The monovalent system (i.e., LgBiT bound to SD36 and SmBiT to HSB2.A or vice versa) showed only a very minor increase in luminescence intensity, which can be explained by low the affinity of the monovalent HSB2.A interaction (**Figure 2E**). Consistent with this, a combination of trivalent HSB2.A fused to LB and trivalent SD36 fused to SB resulted in a much higher dynamic range (285-fold) and better sensitivity (LoD <200 pM) than the corresponding monovalent setup (**Figure 2F**). The latter result demonstrates the advantage of sensor formats that allow trivalent interactions for low-affinity binders such as HSB2.A.

### GLOVID, a generic sensor platform for viral protein detection

Having established the performance of the GLOVID platform for the detection of the IAV HA-trimer, we next explored the modularity of the platform for the detection of several other viral proteins from IAV and RSV using different types of binding domains. First, we tested two other high-affinity nanobody binders: SD38, which binds to the stem region of HA trimers of a different IAV group – such as A/Solomon Islands/3/2006 (H1Sol) ^21^ – and F-VHH-4, which binds to the RSV-F trimer protein in its prefusion conformation ^24^. In both cases, monovalent sensor formats showed very high dynamic ranges (300-1600-fold increase in bioluminescence intensity between 1 pM and 10 nM of target), with LoDs in the low pM range (**Figure 3A**). As expected and also observed for the SD36-based detection of H3HK, sensor formats using trivalent binders proved much less effective (Figure S2). These results show that – if high affinity nanobodies are used – GLOVID can be easily adapted to target viral proteins of different virus strains without compromising sensitivity.

**Figure 3.**
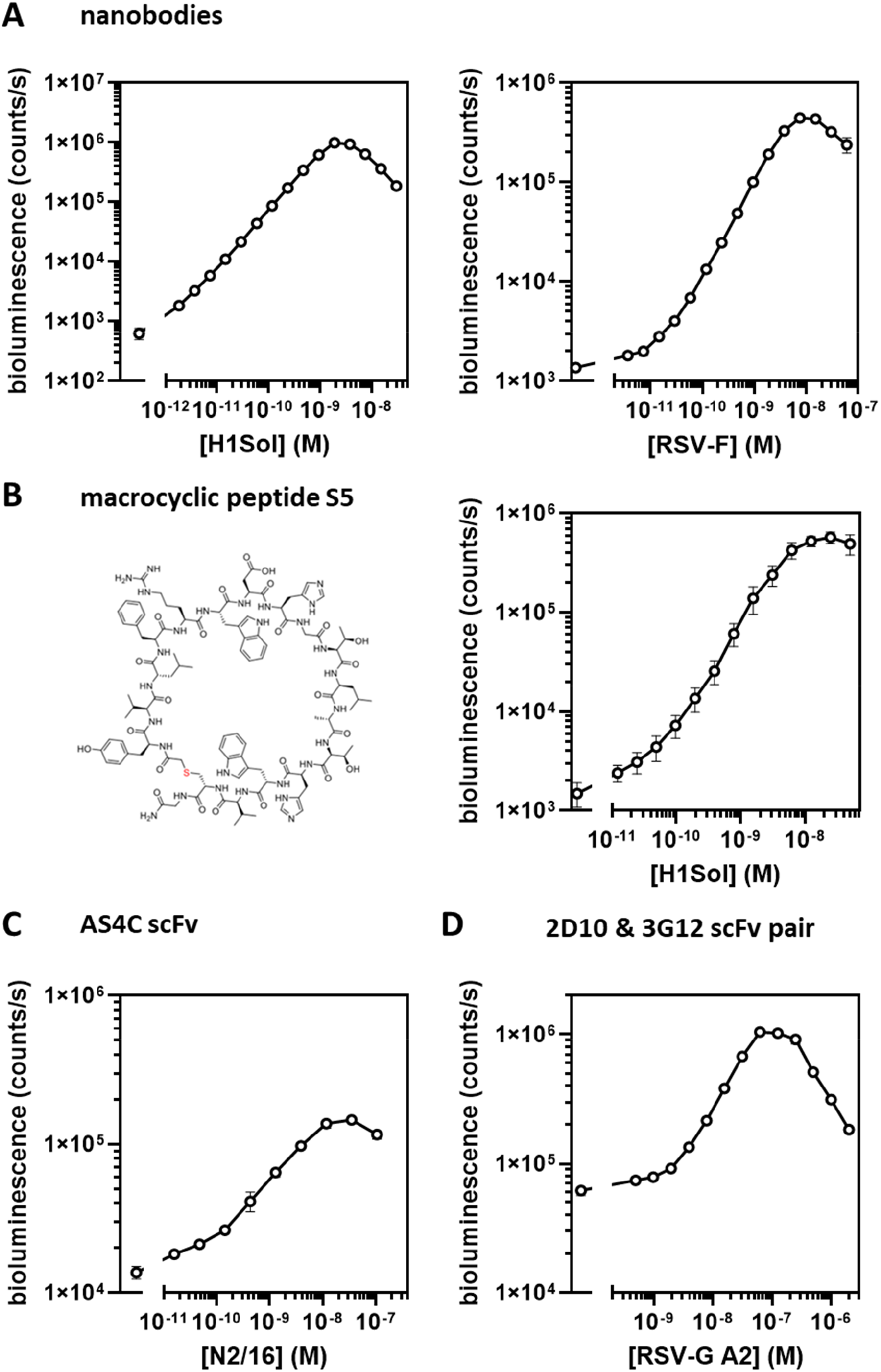
GLOVID assays with different binder classes. A) Assays with nanobody-functionalized monovalent sensor components, targeting IAV H1Sol using SD38 (left) and targeting RSV-F using F-VHH-4 (right). Experimental conditions: 1 nM GLOVID components, 1xPBS + 1mg/ml BSA, final NanoGlo dilution 1:2000, 1 h incubation at 22 °C. LoD RSV mono: 3 pM, DR = 326. LoD = 2 pM, SD38: DR = 1592. B) Assay with monovalent sensor components functionalized with macrocyclic peptide S5, targeting IAV H1Sol (2 nM LgBiT, 6 nM SmBiT, 1xPBS, 1 h incubation at 22 °C, 1:1000 diluted NanoGlo). C) & D) Assays with scFv-functionalized monovalent sensor components, targeting IAV N2/16 using AS4C (c, final GLOVID component concentration 2 nM, 1xPBS + 1 mg/ml BSA, 16 h incubation at 4 °C, final NanoGl dilution 1:2000) and RSV-G A2 (D, 16 nM 2D10-SmBiT, 4 nM 3G12-LgBiT, other conditions as in C)) Error bars correspond to the standard deviation of n=3 replicates. Lines connecting the points are added for better visualisation.

The DogCatcher/DogTag ligation system also offers the opportunity to use binders that cannot be produced using recombinant expression systems. In recent years, several high-throughput display systems have been developed that allow the screening of peptides incorporating non-natural amino acids and non-natural peptide backbones. Using such systems, Pascha and coworkers reported the development of the high-affinity peptide macrocycle S5 (*K*_*D*_ = 5 nM) that inhibits H1 and H5 IAV infection by binding to a conserved epitope in the stem region of HA in its prefusion conformation ^25^. To allow the use of S5 with the GLOVID platform, we used automated Fmoc solid-phase peptide synthesis (SPPS) to synthesize S5 and DogTag peptides containing azide and alkyne functionalities, respectively. The S5 and DogTag peptides were coupled *via* a copper-catalyzed cycloaddition reaction, purified via HPLC and subsequently ligated to LgBiT-Dog1 and SmBiT-Dog1 proteins. Titration experiments with the H1 hemagglutinin trimer protein (H1Sol) showed a robust, 380-fold increase in bioluminescence intensity between 1 pM and 10 nM H1Sol (**Figure 3B**), similar to the results obtained with nanobody binders. As expected ^25^, the GLOVID sensor based on S5 was specific for H1, as no increase in bioluminescence was observed upon titration with the H3-derived H3HK protein (Figure S3).

A third class of important binding domains are single-chain variable fragments (scFv), which, unlike full-sized antibodies, can be recombinantly expressed in *E. coli* or yeast expression systems. Here we created Dog-tagged scFvs derived from antibodies binding to the tetrameric IAV neuraminidase protein (NA) and antibodies binding to different epitopes of the monomeric RSV-G glycoprotein. For targeting tetrameric IAV NA, we selected 1GO1 ^26^ due to its broad NA target binding and AS4C as an N2-specific NA-binding antibody ^23^. For anti-RSV-G binders, we chose antibody templates 3G12 ^27^ and 2D10 ^28^ as they were described to target slightly different RSV-G epitopes. This was considered necessary for binder characterization as soluble RSV-G is monomeric – in contrast to membrane-bound RSV-G which is oligomeric ^29^ – and two non-overlapping binding sites are required for NanoBiT complementation. All scFv designs were fused to DogTag and expressed in yeast (*Pichia pastoris*) to enable the correct formation of disulfide bonds. Microscale thermophoresis (MST) experiments confirmed the binding of purified scFvs to their targets with *K*_*D*_s in the nM range (Figure S4).

After successful conjugation of the scFv-DogTag fusion proteins to LgBiT-Dog1 and SmBiT-Dog1, various titration experiments were performed using NA-tetramer and RSV-G monomer as target proteins. While functional sensors were obtained in all cases (**Figure 3C & D**, Figure S5), the sensor performance of many of the scFv-derived GLOVID sensors was less impressive than that of the nanobody and peptide-derived sensors. The monovalent GLOVID assay that used 1GO1 scFv to detect tetrameric N1/09 (A/CA/07/2009) showed a 39-fold increase in bioluminescence but a low overall bioluminescent signal, indicating that a large portion of conjugated 1GO1 was inactive (Figure S5). Titration experiments with AS4C-LB and AS4C-SB with tetrameric NA (N2/16 (A/NL/354/16)) showed a high bioluminescent signal but also a high background, resulting in a relatively low dynamic range of ∼10-fold (**Figure 3C**). A similar high background luminescence was observed for an assay that combined 3G12-LgBiT with 2D10-SmBiT to detect RSV-G A2, which resulted in a relatively modest 16-fold dynamic range and a LoD of 2 nM (**Figure 3D)**. A possible explanation for the high background could be the formation of scFv hetero-oligomers, where even minor amounts of such oligomers would result in a high background signal. The performance of the 3G12-LgBiT/2D10-SmBiT pair was also found to strongly depend on the affinity of the scFvs, where 3G12 and 2D10 scFvs showed higher affinities towards RSV-G from strain A2 (*K*_*D*, app_ = 8 and 17 nM) than RSV-G from strain B1 (*K*_*D*,app_ = 28 and 300 nM, Figure S4). When RSV-G B1 was used as the target no increase in bioluminescence was observed up to 6.2 µM of RSV-G B1 (Figure S6). Unexpectedly, a control assay that used only 2D10 as binder (2D10-LgBiT/2D10-SmBiT) still showed a small increase in bioluminescence at high concentrations of RSV-G from strain A2 (LoD = 8 nM, DR 6-fold), suggesting the presence of some RSV-G oligomers (Figure S7) ^29^.

## GLOVID detection of intact Influenza A virus

The success of GLOVID in detecting viral surface proteins with high sensitivity encouraged us to explore the potential of the platform for specific detection of intact respiratory virus particles. To avoid detection bias by the presence of ‘free’ surface proteins, we rendered luciferase complementation dependent on the presence of two different viral surface proteins as present on the surface of intact virions (**Figure 1A**). H1N2 swine influenza virus (A/swine/Italy/150383-1/2014, provided by the European Virus Archive) was chosen as a target, using the monovalent S5 peptide to target HA (fused to LB-Dog1) and the AS4C scFv (fused to SB-Dog1 and SB-Dog3) to target the N2 NA. The number of viral genomes in this preparation was determined by quantitative reverse-transcription droplet digital PCR (RT-ddPCR) targeting the M gene of H1N2, which revealed ∼4.6· 10^8^ RNA copies per ml stock solution. Viral RNA in the viral stock is considered stable inside the virions, whereas viral transcripts (mRNA) and transcription intermediates (cRNA) are prone to degradation by RNases present in the cell lysate ^30,31^. Considering that the majority of intact virions contains a single copy of the M gene^31^, in this case the concentration of genomic units of 760 fM is likely to be similar to the amount of virus particles.

Titration experiments showed a 20-fold increase in bioluminescent intensity with a very low LoD of 0.14 fM of genomic units (5σ criterium) (**Figure 4A**), which corresponds to a RT-qPCR Ct value of ∼29 (Figure S8). Surprisingly, no further increase in bioluminescence was observed above a concentration of 0.5 fM genomic units. As a control, a GLOVID assay with a combination of the two H1 binders SD38 and S5 was used, which resulted in a similar increase in bioluminescent signal up until ≈ 0.4 – 0.6 fM; however, in this case the signal increased further with increasing virus concentration (**Figure 4B**). The difference in the sensor response observed for higher virion concentrations is puzzling. It suggests that complex formation involving different surface proteins is more sensitive to sensor dilution than complex formation targeting homotrimeric proteins, possibly because of higher effective concentrations in the latter. Control titration experiments using 1:1 mixtures of free HA and NA did not show an increase in bioluminescence, proving that complementation of the S5/AS4C pair occurs exclusively on the virion surface and is not sensitive to the presence of free protein (**Figure 4C**). The LoD of 0.14 fM for direct, amplification-free in-solution detection of intact H1N2 virions is lower than the average RNA load of respiratory virus in oral fluid (7 × 10^6^ copies/ml or 11.6 fM, ^32^) and consistent with the 10^6^ copies/mL (1.6 fM) required for established IAV antigen tests ^33^. The ability to detect such low concentrations of virus even at relatively high sensor concentrations (2-6 nM) is enabled by the presence of multiple copies of target protein per virion (e.g. ≈300-500 HAs ^34^) and the high dynamic range (low background) of the GLOVID assay (Figure S1).

**Figure 4.**
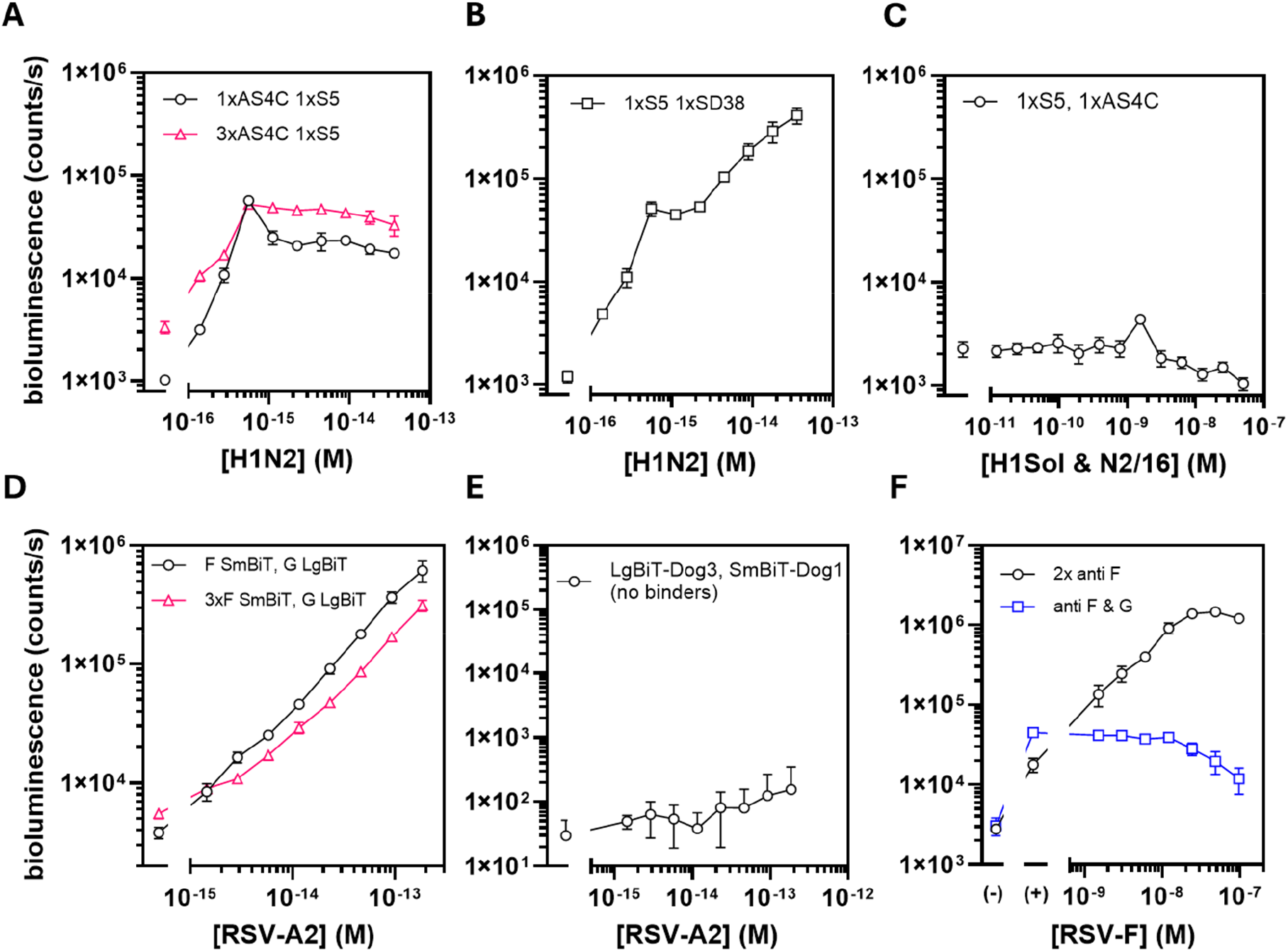
Direct detection of intact virions using GLOVID. A) GLOVID assay targeting H1N2 via S5-LgBiT and monovalent AS4C-SmBiT (black circles), and S5-LgBiT and trivalent AS4C-SmBiT (pink triangles). B) GLOVID assay targeting H1N2 via SD38-LgBiT and S5-SmBiT (both HA specific). Experimental conditions in A) and B): 2 nM LgBiT, 6 nM SmBiT, 1xPBS, final NanoGlo dilution 1:2000, 1 h incubation at 22 °C. C) Response of an assay containing 2 nM S5-LgBiT and 6 nM AS4C-SmBiT to an equimolar mixture of H1Sol and N2/16 proteins. Experimental conditions: 1xPBS, incubation 1 h 22 °C, 1:2000 diluted NanoGlo substrate. D) GLOVID assays targeting RSV-A2 via 2D10-LgBiT / F-VHH-4-SmBiT (monovalent or trivalent). Experimental conditions: 4 nM final GLOVID component concentration, 1xPBS, final NanoGlo dilution 1:2000, 2 h incubation 22 °C. E) Control experiment for GLOVID on RSV A2 virus sample where no binders were conjugated to LgBiT-Dog3 and SmBiT-Dog1. Experimental conditions: 4 nM LgBiT-Dog3 and SmBiT-Dog1, 1xPBS, final NanoGlo dilution 1:1000, 2 h incubation at 22 °C. F) Experiment in which additional RSV-F was spiked into samples that contained 37 fM RSV-A2, followed by testing with GLOVID assemblies F-VHH-4-LgBiT / F-VHH-4-SmBiT (black circles) or 2D10-LgBiT / F-VHH-4-SmBiT (blue squares). (-) refers to the buffer control, (+) refers to 37 fM of added RSV A2 particles as the basis for the spiking experiment. Experimental conditions: 2 nM LgBiT, 6 nM SmBiT, 1xPBS, final NanoGlo dilution 1:1000, 1 h incubation at 22 °C. Error bars represent the standard deviation of n=3 replicates. Lines are used to connect data points for better visualisation. Please note that all concentrations in A, B, D and E are based on RT-ddPCR and reflect the concentration of genomic units, which can be either higher or lower than the concentration of virus particles.

### GLOVID detection of respiratory syncytial virus particles

To establish the general applicability of the GLOVID platform, we also tested its performance on a laboratory strain of RSV, RSV-A2 ^35^. The concentration of matrix protein-coding RNA in the stock, as assessed via RT-ddPCR, was determined to be 6.7 ·10^9^ cp per ml, corresponding to a concentration of 11 pM of genomic units. CryoTEM studies have shown that RSV virus particles can vary substantially in shape and size, and typically contain multiple genomic copies (between 1 and 9), which means that the number of virus particles is probably lower than the amount of genomic units ^36^. The infectivity of the RSV-A2 stock preparation was also assessed in a plaque staining assay on Vero cells, showing a concentration of 8.1·10^8^ plaque forming units (PFU) per ml, which is 12% of the amount of genomic units. However, amount of plaque forming units can also underestimate the amount of virus particles. We therefore decided to use the concentration of genomic units in our titration experiments.

As with IAV, purified RSV-A2 samples may contain ‘free’, soluble viral proteins, so we targeted two distinct surface proteins to make the bioluminescent signal dependent on the presence of intact RSV particles. Therefore, anti-RSV-G scFv 2D10 was conjugated to LgBiT-Dog1, and prefusion-RSV-F-specific F-VHH-4 was conjugated to SmBiT-Dog1 and SmBiT-Dog3. Titrations with RSV-A2 showed an increase in bioluminescent signal with increasing virus concentration for both sensor configurations, resulting in LoDs of 1.4 – 3 fM (5σ criterium) **(Figure 4D)**, which corresponds to a Ct value of ∼23 (Figure S8). A control experiment with no binders attached to the GLOVID components showed no significant increase in signal above the background (**Figure 4E**). Titration experiments where either only RSV-F or only RSV-G were targeted gave similar results as for the heterobivalent assay, with LoDs of ∼1.4 – 3 fM of genomic units for the F-VHH-4 based assay and 1.2 fM of genomic units for an 2D10- and 3G12-based, monovalent sandwich assay (Figure S9). The latter results suggest that the used RSV-A2 stock contained little to no impurities of free surface proteins but mainly intact virions. To confirm that the heterobivalent GLOVID is insensitive to the presence of free surface proteins, we also titrated purified, prefusion-stabilized RSV-F into samples with a constant amount of RSV-A2 (37 fM of genomic units). An additional signal increase was only observed in case both sensor components targeted RSV-F, and not in case RSV-F and RSV-G were targeted (**Figure 4F**). In fact, at high concentrations of spiked RSV-F protein, the signal from the heterobivalent GLOVID decreased slightly, in line with free RSV-F outcompeting RSV-A2 for F-VHH-4 binding.

We also explored the possibility of using GLOVID to detect RSV in clinical nasal swab samples taken from patients, but preliminary experiments were unsuccessful. To check if the matrix complexity caused issues, we spiked purified RSV-A2 virions into different matrices, including FastAmp (Intact Genomics) and an RSV-negative nasal swab sample diluted to 15% (Figure S10). While the GLOVID assay showed an increase in bioluminescence upon the addition of RSV-A2 to an RSV-negative nasal swab sample, the assay’s sensitivity was found to be considerably reduced, mainly due to a higher background signal. In addition, the anti-RSV binders used in our assay may have a lower affinity for currently circulating RSV strains than for the laboratory strain RSV-A2, which would further lower the sensitivity of the GLOVID assay.

## Discussion

In this study, we developed a modular bioluminescent sensor platform for intact virion detection using the DogTag/DogCatcher system to fuse split luciferase domains to a wide variety of binding moieties. We confirmed that the reaction between DogCatcher and DogTag is spontaneous and efficient, proceeding to apparent completion irrespective of the binder attached to the DogTag ^14^. Using high-affinity binders, purified viral surface proteins were detected with LoDs in the low pM range. The trivalent display of the low-affinity, computationally designed H3HK head binder HSB2.A rendered it ‘functional’ and allowed detection of H3HK by combining it in a sandwich with the high-affinity nanobody binder SD36. Finally, ultrasensitive detection of intact H1N2 IAV was achieved against a background of ‘free’ HA proteins by combining the anti-H1 macrocyclic peptide S5 with the anti-N2 scFv AS4C created from an existing IgG antibody (**Figure 4A**). Intact RSV-A2 detection with anti-F protein and anti-G protein binders was successful for all combinations of binders (**Figure 4C**, Figure S9).

GLOVID enables fast and ultrasensitive quantification of respiratory viruses by directly detecting intact virions at a limit of detection in the low fM range. The GLOVID platform thus offers distinct advantages compared to current homogeneous, bioluminescent assays that directly detect viral proteins. The analytical performance of GLOVID is similar to that of RAPPID ^12,37^, but GLOVID allows a much broader range of binders and enables affinity enhancement through multivalent interactions. In a recent study, an unusual pseudo-luciferase activity of SARS-CoV-2 S trimer that leads to oxidation of *Cypridina* luciferin allowed swift and accurate detection of the viral surface protein. However, the detection limit was only ∼2 nM and the detection mechanism is likely to be restricted to SARS-CoV-2 ^38^. Hunt and coworkers recently reported a single protein sensor (FUS231-P12) that uses trivalent binding to the SARS-CoV-2 S trimer to increase BRET between a luciferase and an acceptor fluorescent domain. This sensor was sensitive with a LoD of 11 pM ^39^, but in its current form was not specific to intact virus. Our group recently reported a similar type of single protein sensors where multivalent binding of the sensor to trivalent viral surface proteins induces a conformational switch that allows intramolecular complementation of split NanoLuc ^23^. However, even the most optimized versions of these Dark-LUX sensor proteins were 10-fold less sensitive compared to the corresponding GLOVID sensor when targeting the same surface protein and using the same binding domains (e.g., 3.7 pM vs. 40 pM for H3HK detection) ^23^. The Dark-LUX sensors also did not distinguish between free viral proteins and intact virus.

Although our first effort to detect RSV in clinical samples failed, we anticipate that this can be remedied by using high-affinity binders for currently circulating virus strains, or by including a low affinity auto-inhibitor domain in the sensor design to prevent high background signal ^40^. Notably, the modularity of the GLOVID platform allows straightforward screening for optimal binder pairs with high affinity and low background interactions. More elaborate protein architectures based on both DogCatcher/DogTag and SpyCatcher/SpyTag variants can also be envisioned ^41^, e.g. by fusing one of the NanoBiT domains to multiple binders that target different epitopes to achieve increased target affinity and selectivity.

While GLOVID can distinguish virions from free proteins, it does not necessarily inform if the detected virions are infectious. The correlation between virus amount and infectivity is highly complex and varies between different viruses and strains ^42,43^, as we also observed in this study for RSV when comparing the ddPCR and PFU/ml values. Infectivity may also depend on surface protein conformations (e.g. RSV-F with pre- and post-fusion state). It therefore remains to be established whether the GLOVID assay is a good proxy for infectivity, and if so, how this depends on the type of virus. Nonetheless, we believe that GLOVID may provide an attractive future alternative for the laborious cell-based assays that are currently used to assess the infectivity of (pseudo-) viruses ^44,45^.

## Material and Methods

### Cloning

DNA nucleotides and gblocks or gene fragments were purchased from IDT or Twist Biosciences, and both cloning and plasmid amplification was performed in *Escherichia coli* TOP10. gblocks or gene fragments were inserted into pET28a(+) vectors via restriction and ligation with T4 Ligase (NEB). Specifically, for construction of trivalent GLOVID scaffolds LgBiT-3x-DogCatcher and 3xDogCatcher-SmBiT, three separate DogCatcher gene fragments – two of which contained (TP)_21_ linkers with scrambled codons ^46^ were ligated to pET28a(+) containing LgBiT and Strep-tag II or SmBiT110 and 6xHis-Tag, respectively.

Construction of plasmids containing SD36-DogTag, SD38-DogTag, HSB2.A-DogTag and AS4C-HL-DogTag was described previously ^23^. All scFvs were designed in *heavy chain-light chain* configuration (HL) and cloned to pPICZalphaB using pPICZalphaB-SapL3 (addgene plasmid #78171) as template. Genomic integration of scFv constructs into *P. pastoris* and subsequent purification was performed as described previously for AS4C-HL-DogTag ^23^.

### Protein purification of genetically encoded GLOVID components

HSB2.A and GLOVID scaffold with DogCatchers were produced in *E. coli* BL21 (DE3), whereas nanobodies (VHHs) were produced in *E. coli* Shuffle® T7 (NEB). The bacteria were transformed with the corresponding pET28a(+) plasmids and grown in LB medium supplemented with 50 µg/ml kanamycin at 37 °C at 180 rpm in a shaking incubator. Large cultures (0.5-2 l) in 2 l or 5 l baffled flasks were inoculated with corresponding overnight cultures and induced with IPTG at OD_600_ 0.6-0.8. Proteins were expressed overnight at 18 °C. Harvested cells were lysed with BugBuster reagent (Novagen) supplemented with Benzonase and cOmplete™ protease inhibitor tablets, and proteins were purified according to their tags (see protein sequences) with Ni-NTA chromatography and/or Strep-Tactin XT (IBA) using gravity flow columns. For SD36-DogTag and SD38-DogTag, an additional size exclusion step was performed as described in ^23^. Protein purity was confirmed by reducing SDS-PAGE and concentrations were calculated using A_280_ Nanodrop measurements using the corresponding theoretical extinction coefficients (based on protein sequence). Proteins were aliquoted, flash frozen in liquid N_2_ and stored at -70 °C.

### Protein purification of viral surface proteins

RSV-G A2 and RSV-G B1 proteins were purchased from Sino Biological. Hemagglutinins H1Sol and H3HK from A/Solomon Islands/3/2006 (H1N1) and A/Hong Kong/1/1968 (H3N2), and neuraminidase N2/16 (A/NL/354/16) were purified from HEK293 cell culture supernatants as described previously ^20,23^. N1/09 (A/CA/07/2009) was a kind gift from Pramila Rijal and Alan Townsend ^47^.

### Synthesis and purification of macrocyclic peptide S5

Peptides were prepared using standard Fmoc solid phase peptide synthesis on a Chorus synthesizer (Gyros protein technologies, Sweden) on tentagel XV RAM resin (Iris biotechnology, Germany) at 25 µmol scale. Resin was swelled in 3 ml dimethyl formamide (DMF) for 30 min, then coupling cycles were carried out with cycles of deprotection (20% piperidine with 0.1 M oxyma, 3 ml, at 80 °C for 90 s), washing (three times, 3 ml DMF), coupling (5 eq. amino acid with 10 eq. DIC and 5 eq. oxyma, 3 ml, at 55 °C for 15 min), capping (2 M acetic anhydride and 2 M pyridine in DMF, 3 ml, 5 min at room temp), and washing (three times, 3 ml DMF). No final deprotection was carried out. After synthesis, resin was washed with dichloromethane (three times, 3 ml each) and then dried by vacuum for 30 min. Dried resin was then swelled in 1.5 ml cleavage solution (90% trifluoroacetic acid with 5% water, 2.5% triisopropyl silane, 2.5% 2,2 -(ethylenedioxy)diethanethiol as scavengers) and incubated at room temperature with gentle shaking for 3 hours. Peptide was precipitated by addition of cleavage solution into 30 ml ice cold diethyl ether, which was then pelleted by centrifugation at 5,000 rcf for 5 min. The pellet was washed by resuspending in 10 ml diethyl ether and re-pelleting as before for a total of 3 cycles. The final pellet was then air dried and dissolved in 2 ml dimethyl sulfoxide. HA S5 peptide was cyclized in solution by addition of *circa* 250 µl of *N,N*-diisopropylethylamine (5 drops) and allowing to stand at room temperature until LC-MS analysis indicated reaction was complete. The reaction was then quenched by addition of *circa* 250 µl (5 drops) of trifluoroacetic acid, and both peptides were purified by RP-HPLC (C18 column, elution gradient from 10-70% v/v acetonitrile in water with 0.1% trifluoroacetic acid).

Sequence for HA S5 ^25^ GGS azide:

(Chloroacetic acid)-YVLFRWDHGTLATHWVCGGSGGS-(azidohomoalanine)-G-NH_2_

Sequence for alkyne dog tag:

(4-Pentynoic acid)-GGDIPATYEFTDGKHYITNEPIPPKG-NH_2_

Click conjugation of the two peptides was carried out at 500 nmol scale, with a slight excess of S5 peptide (650 nmol, 1.3 eq.). CuSO_4_ (5 µmol, 10 eq.) was first mixed with THPTA ligand (7.5 µmol, 15 eq.) whereupon the colour became a deep blue, and then reduced to Cu^I^ by addition of sodium ascorbate (25 µmol, 50 eq.) with change in color to pale yellow. To this was then added a pre-mixed solution of Tris buffer at pH 8.5 (250 mM final concentration), dimethyl sulfoxide (final concentration 50% v/v) and each of the two peptides from stock solutions in dimethyl sulfoxide, in a final reaction volume of 75 µl. The reaction was allowed to proceed at room temperature, and then stored at -20 °C after 2.5 hours before final purification again by RP-HPLC after addition of trifluoroacetic acid until acidic (Figure S11, S12). The purified product was freeze dried, and the concentration of a stock solution determined using a calculated extinction coefficient at 280 nm of 15470 M^-1^cm^-1^.

### Virus strains

A preparation of Swine influenza A/swine/Italy/114922/2014 (H1N2) with an HA titer of 64 was obtained via the European Virus Archive Global (EVAg) https://www.european-virus-archive.com/virus/swine-influenza-aswineitaly1149222014-h1n2. The RSV-A2 stock, an A subtype of RSV (ATCC, VR-1540, Rockville) was prepared as previously described ^24^. Briefly, the virus was propagated on HEp-2 cells, purified using PEG precipitation and quantified on Vero cells by plaque assay using goat anti-RSV serum (AB1128, Chemicon International). No unexpected or unusually high safety hazards were encountered.

### Sensor assembly

DogCatcher/DogTag ligations for monovalent GLOVID variants was performed by mixing DogCatcher parts and DogTag parts at equimolar concentration, at least at 0.75 µM, usually at 5 µM each. Protein ligations for trivalent variants were performed at a molar ratio of 1:4 to guarantee complete conversion of all added DogCatcher parts into conjugated sensor components. Each assembly reaction was shortly spun down after mixing, and incubated overnight at room temperature (22 °C) followed by storage at 4 °C. For each GLOVID assay, new assemblies were prepared if the last assemblies were older than 3 days. Reactions were performed in a final volume of 50 µl and with 1xPBS as buffer. Assembly success was routinely confirmed via reducing SDS-PAGE.

## Titrations

Titrations were performed as described previously in ^40^. Unless stated otherwise, titrations were performed in low protein binding tubes in 1xPBS supplemented with 1 mg/ml bovine serum albumin (BSA). NanoGlo substrate (furimazine, Promega) was added from a freshly prepared 10x stock after incubation (1 h at 22 °C unless stated otherwise), directly before the measurement to obtain the final volume and concentrations. Unless stated otherwise, the final NanoGlo dilution during the measurements was 1:2000. Assays were performed in white 364-well plates (flat bottom) and a total volume of 20 µl with n=3 technical replicates. Unless stated otherwise, shown bioluminescence values were determined from total bioluminescence (integration time 100 ms) measured at 22 °C in a TECAN Spark 10M. The dynamic range (DR) of each measurement was calculated via

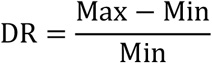

with *Max* being the signal at saturated sensor (mean of n=3) and *Min* the signal in absence of the target (mean of n=3). DR uncertainties were propagated from the standard deviation of *Max* and *Min* (*s(Max)* and *s(Min)*, respectively, using ^48^

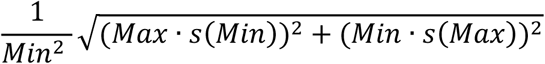

The limit of detection (LoD) of each sensor was calculated by applying the 5σ rule (mean of blank plus 5 times its standard deviation) similar to previous studies ^40^.

## Acknowledgments

The authors thank the group of Bruno Correia for providing purified, prefusion-stabilized RSV-F. We would like to thank Pramila Rijal and Alain Townsend for kindly providing N1/09 and the sequence of the N2-specific AS4C antibody. We would like to thank the members of the Merkx group for fruitful discussions and feedback.

## Author contributions

AG conceived and designed the study, performed experiments, analyzed the data, and wrote the manuscript. HvdV performed ddPCR experiments and provided important feedback. SAKJ synthesized and purified the DogTagged macrocyclic peptide S5 and provided important feedback. JF collected RSV samples and provided important feedback. IR purified and provided RSV-A2 and initially characterized F-VHH-4, RPdV provided viral surface proteins and important feedback, XS provided RSV-A2 and important feedback. MM supervised the study, designed experiments, analyzed the data, and wrote the manuscript. All authors discussed the results and commented on the manuscript.

## Ethical statement

This study was performed in line with the principles of the Declaration of Helsinki and in accordance with national regulation. Work with anonymized residual material from RSV patients was approved prior to the experiments by the Ethical Review Board of Eindhoven University of Technology and the Rijnstate Hospital, Arnhem, The Netherlands.

## Data and materials availability

The authors will share all raw data produced in this study upon request.

## Supporting Information

Additional materials and methods, supporting figures and protein sequences (PDF)

## References

(1) Salgado, C. D.; Farr, B. M.; Hall, K. K.; Hayden, F. G. Influenza in the Acute Hospital Setting. Lancet Infect Dis 2002, 2 (3), 145–155. 10.1016/S1473-3099(02)00221-9.

(2) Hill-Ricciuti, A.; Walsh, E. E.; Greendyke, W. G.; Choi, Y.; Barrett, A.; Alba, L.; Branche, A. R.; Falsey, A. R.; Phillips, M.; Finelli, L.; Saiman, L. Clinical Impact of Healthcare-Associated Respiratory Syncytial Virus in Hospitalized Adults. Infect Control Hosp Epidemiol 2023, 44 (3), 433–439. 10.1017/ice.2022.128.

(3) Bruning, A. H. L.; Leeflang, M. M. G.; Vos, J. M. B. W.; Spijker, R.; de Jong, M. D.; Wolthers, K. C.; Pajkrt, D. Rapid Tests for Influenza, Respiratory Syncytial Virus, and Other Respiratory Viruses: A Systematic Review and Meta-Analysis. Clinical Infectious Diseases 2017, 65 (6), 1026–1032. 10.1093/cid/cix461.

(4) Song, Q.; Sun, X.; Dai, Z.; Gao, Y.; Gong, X.; Zhou, B.; Wu, J.; Wen, W. Point-of-Care Testing Detection Methods for COVID-19. Lab Chip 2021, 21 (9), 1634–1660. 10.1039/D0LC01156H.

(5) Laue, M.; Hoffmann, T.; Michel, J.; Nitsche, A. Visualization of SARS-CoV-2 Particles in Naso/Oropharyngeal Swabs by Thin Section Electron Microscopy. Virol J 2023, 20 (1), 21. 10.1186/s12985-023-01981-9.

(6) Klasse, P. J. Molecular Determinants of the Ratio of Inert to Infectious Virus Particles. In Progress in Molecular Biology and Translational Science; Klasse, P. J., Ed.; 2015; Vol. 129, pp 285–326. 10.1016/bs.pmbts.2014.10.012.

(7) Heider, S.; Metzner, C. Quantitative Real-Time Single Particle Analysis of Virions. Virology 2014, 462–463, 199–206. 10.1016/j.virol.2014.06.005.

(8) Wang, S.; Zhang, F.; Mei, M.; Wang, T.; Yun, Y.; Yang, S.; Zhang, G.; Yi, L. A Split Protease-E. Coli ClpXP System Quantifies Protein–Protein Interactions in Escherichia Coli Cells. Commun Biol 2021, 4 (1), 841. 10.1038/s42003-021-02374-w.

(9) Bae, J.; Kim, J.; Choi, J.; Lee, H.; Koh, M. Split Proteins and Reassembly Modules for Biological Applications. ChemBioChem 2024, 25 (10). 10.1002/cbic.202400123.

(10) van der Veer, H. J.; van Aalen, E. A.; Michielsen, C. M. S.; Hanckmann, E. T. L.; Deckers, J.; van Borren, M. M. G. J.; Flipse, J.; Loonen, A. J. M.; Schoeber, J. P. H.; Merkx, M. Glow-in-the-Dark Infectious Disease Diagnostics Using CRISPR-Cas9-Based Split Luciferase Complementation. ACS Cent Sci 2023, 9 (4), 657– 667. 10.1021/acscentsci.2c01467.

(11) de Stigter, Y.; van der Veer, H. J.; Rosier, B. J. H. M.; Merkx, M. Bioluminescent Intercalating Dyes for Ratiometric Nucleic Acid Detection. ACS Chem Biol 2024, 19 (2), 575–583. 10.1021/acschembio.3c00755.

(12) Ni, Y.; Rosier, B. J. H. M.; van Aalen, E. A.; Hanckmann, E. T. L.; Biewenga, L.; Pistikou, A. M. M.; Timmermans, B.; Vu, C.; Roos, S.; Arts, R.; Li, W.; de Greef, T. F. A.; van Borren, M. M. G. J.; van Kuppeveld, F. J. M.; Bosch, B. J.; Merkx, M. A Plug-and-Play Platform of Ratiometric Bioluminescent Sensors for Homogeneous Immunoassays. Nat Commun 2021, 12 (1). 10.1038/s41467-021-24874-3.

(13) Dixon, A. S.; Schwinn, M. K.; Hall, M. P.; Zimmerman, K.; Otto, P.; Lubben, T. H.; Butler, B. L.; Binkowski, B. F.; MacHleidt, T.; Kirkland, T. A.; Wood, M. G.; Eggers, C. T.; Encell, L. P.; Wood, K. V. NanoLuc Complementation Reporter Optimized for Accurate Measurement of Protein Interactions in Cells. ACS Chem Biol 2016, 11 (2), 400–408. 10.1021/acschembio.5b00753.

(14) Keeble, A. H.; Yadav, V. K.; Ferla, M. P.; Bauer, C. C.; Chuntharpursat-Bon, E.; Huang, J.; Bon, R. S.; Howarth, M. DogCatcher Allows Loop-Friendly Protein-Protein Ligation. Cell Chem Biol 2022, 29 (2), 339-350.e10. 10.1016/j.chembiol.2021.07.005.

(15) Rossey, I.; Hsieh, C.-L.; Sedeyn, K.; Ballegeer, M.; Schepens, B.; McLellan, J. S.; Saelens, X. A Vulnerable, Membrane-Proximal Site in Human Respiratory Syncytial Virus F Revealed by a Prefusion-Specific Single-Domain Antibody. J Virol 2021, 95 (11). 10.1128/jvi.02279-20.

(16) Wilson, R. H.; Morton, S. K.; Deiderick, H.; Gerth, M. L.; Paul, H. A.; Gerber, I.; Patel, A.; Ellington, A. D.; Hunicke-Smith, S. P.; Patrick, W. M. Engineered DNA Ligases with Improved Activities in Vitro. Protein Engineering, Design and Selection 2013, 26 (7), 471–478. 10.1093/PROTEIN/GZT024.

(17) Gräwe, A.; Merkx, M.; Stein, V. IFLinkC-X: A Scalable Framework to Assemble Bespoke Genetically Encoded Co-Polymeric Linkers of Variable Lengths and Amino Acid Composition. Bioconjug Chem 2022, 33 (7), 1415–1421. 10.1021/acs.bioconjchem.2c00250.

(18) Sørensen, C. S.; Jendroszek, A.; Kjaergaard, M. Linker Dependence of Avidity in Multivalent Interactions Between Disordered Proteins. J Mol Biol 2019, 431 (24), 4784–4795. 10.1016/j.jmb.2019.09.001.

(19) Gräwe, A.; Stein, V. Linker Engineering in the Context of Synthetic Protein Switches and Sensors. Trends Biotechnol 2021, 39 (7), 731–744. 10.1016/J.TIBTECH.2020.11.007.

(20) van der Woude, R.; Turner, H. L.; Tomris, I.; Bouwman, K. M.; Ward, A. B.; de Vries, R. P. Drivers of Recombinant Soluble Influenza A Virus Hemagglutinin and Neuraminidase Expression in Mammalian Cells. Protein Science 2020, 29 (9), 1975–1982. 10.1002/pro.3918.

(21) Laursen, N. S.; Friesen, R. H. E.; Zhu, X.; Jongeneelen, M.; Blokland, S.; Vermond, J.; Van Eijgen, A.; Tang, C.; Van Diepen, H.; Obmolova, G.; Van Der Neut Kolfschoten, M.; Zuijdgeest, D.; Straetemans, R.; Hoffman, R. M. B.; Nieusma, T.; Pallesen, J.; Turner, H. L.; Bernard, S. M.; Ward, A. B.; Luo, J.; Poon, L. L. M.; Tretiakova, A. P.; Wilson, J. M.; Limberis, M. P.; Vogels, R.; Brandenburg, B.; Kolkman, J. A.; Wilson, A. Universal Protection against Influenza Infection by a Multidomain Antibody to Influenza Hemagglutinin. Science (1979) 2018, 362 (6414), 598–602. 10.1126/science.aaq0620.

(22) Strauch, E. M.; Bernard, S. M.; La, D.; Bohn, A. J.; Lee, P. S.; Anderson, C. E.; Nieusma, T.; Holstein, C. A.; Garcia, N. K.; Hooper, K. A.; Ravichandran, R.; Nelson, J. W.; Sheffler, W.; Bloom, J. D.; Lee, K. K.; Ward, A. B.; Yager, P.; Fuller, D. H.; Wilson, I. A.; Baker, D. Computational Design of Trimeric Influenza-Neutralizing Proteins Targeting the Hemagglutinin Receptor Binding Site. Nat Biotechnol 2017, 35 (7), 667– 671. 10.1038/nbt.3907.

(23) Gräwe, A.; Spruit, C. M.; de Vries, R. P.; Merkx, M. Bioluminescent Detection of Viral Surface Proteins Using Branched Multivalent Protein Switches. RSC Chem Biol 2024, 5 (2), 148–157. 10.1039/D3CB00164D.

(24) Rossey, I.; Gilman, M. S. A.; Kabeche, S. C.; Sedeyn, K.; Wrapp, D.; Kanekiyo, M.; Chen, M.; Mas, V.; Spitaels, J.; Melero, J. A.; Graham, B. S.; Schepens, B.; McLellan, J. S.; Saelens, X. Potent Single-Domain Antibodies That Arrest Respiratory Syncytial Virus Fusion Protein in Its Prefusion State. Nat Commun 2017, 8. 10.1038/ncomms14158.

(25) Pascha, M. N.; Thijssen, V.; Egido, J. E.; Linthorst, M. W.; Van Lanen, J. H.; Van Dongen, D. A. A.; Hopstaken, A. J. P.; Van Kuppeveld, F. J. M.; Snijder, J.; De Haan, C. A. M.; Jongkees, S. A. K. Inhibition of H1 and H5 Influenza A Virus Entry by Diverse Macrocyclic Peptides Targeting the Hemagglutinin Stem Region. ACS Chem Biol 2022, 17 (9), 2425–2436. 10.1021/acschembio.2c00040.

(26) Stadlbauer, D.; Zhu, X.; McMahon, M.; Turner, J. S.; Wohlbold, T. J.; Schmitz, A. J.; Strohmeier, S.; Yu, W.; Nachbagauer, R.; Mudd, P. A.; Wilson, I. A.; Ellebedy, A. H.; Krammer, F. Broadly Protective Human Antibodies That Target the Active Site of Influenza Virus Neuraminidase. Science (1979) 2019, 366 (6464), 499–504. 10.1126/science.aay0678.

(27) Fedechkin, S. O.; George, N. L.; Nuñez Castrejon, A.M.; Dillen, J. R.; Kauvar, L. M.; DuBois, R. M. Conformational Flexibility in Respiratory Syncytial Virus G Neutralizing Epitopes. J Virol 2020, 94 (6). 10.1128/JVI.01879-19.

(28) Fedechkin, S. O.; George, N. L.; Wolff, J. T.; Kauvar, L. M.; DuBois, R. M. Structures of Respiratory Syncytial Virus G Antigen Bound to Broadly Neutralizing Antibodies. Sci Immunol 2018, 3 (21). 10.1126/sciimmunol.aar3534.

(29) Escribano-Romero, E.; Rawling, J.; García-Barreno, B.; Melero, J. A. The Soluble Form of Human Respiratory Syncytial Virus Attachment Protein Differs from the Membrane-Bound Form in Its Oligomeric State but Is Still Capable of Binding to Cell Surface Proteoglycans. J Virol 2004, 78 (7), 3524–3532. 10.1128/JVI.78.7.3524-3532.2004.

(30) Girard, J.; Jakob, C.; Toews, L. K.; Fuchs, J.; Pohlmann, A.; Franzke, K.; Kolesnikova, L.; Jeney, C.; Beer, M.; Bron, P.; Schwemmle, M.; Bolte, H. Disruption of Influenza Virus Packaging Signals Results in Various Misassembled Genome Complexes. J Virol 2023, 97 (10). 10.1128/jvi.01076-23.

(31) Chou, Y.; Vafabakhsh, R.; Doğanay, S.; Gao, Q.; Ha, T.; Palese, P. One Influenza Virus Particle Packages Eight Unique Viral RNAs as Shown by FISH Analysis. Proceedings of the National Academy of Sciences 2012, 109 (23), 9101–9106. 10.1073/pnas.1206069109.

(32) Wölfel, R.; Corman, V. M.; Guggemos, W.; Seilmaier, M.; Zange, S.; Müller, M. A.; Niemeyer, D.; Jones, T. C.; Vollmar, P.; Rothe, C.; Hoelscher, M.; Bleicker, T.; Brünink, S.; Schneider, J.; Ehmann, R.; Zwirglmaier, K.; Drosten, C.; Wendtner, C. Virological Assessment of Hospitalized Patients with COVID-2019. Nature 2020, 581, 465. 10.1038/s41586-020-2196-x.

(33) Akashi, Y.; Suzuki, H.; Ueda, A.; Hirose, Y.; Hayashi, D.; Imai, H.; Ishikawa, H. Analytical and Clinical Evaluation of a Point-of-Care Molecular Diagnostic System and Its Influenza A/B Assay for Rapid Molecular Detection of the Influenza Virus. Journal of Infection and Chemotherapy 2019, 25 (8), 578–583. 10.1016/j.jiac.2019.02.022.

(34) Otterstrom, J. J.; Brenburg, B.; Koldijk, M. H.; Juraszek, J.; Tang, C.; Mashaghi, S.; Kwaks, T.; Goudsmit, J.; Vogels, R.; Friesen, R. H. E.; Van Oijen, A. M. Relating Influenza Virus Membrane Fusion Kinetics to Stoichiometry of Neutralizing Antibodies at the Single-Particle Level. Proc Natl Acad Sci U S A 2014, 111 (48), E5143–E5148. 10.1073/pnas.1411755111.

(35) Pandya, M.; Callahan, S.; Savchenko, K.; Stobart, C. A Contemporary View of Respiratory Syncytial Virus (RSV) Biology and Strain-Specific Differences. Pathogens 2019, 8 (2), 67. 10.3390/pathogens8020067.

(36) Kiss, G.; Holl, J. M.; Williams, G. M.; Alonas, E:; Vanover, D.; Lifland, A. W.; Gudheti, M.; Guerrero-Ferreira, R. C.; Nair, V.; Yi, H.; Graham, B. S.; Santangelo, P. J.; Wright, E. R. Structural analysis of respiratory syncytial virus reveals the position of M2-1 between the matrix protein and the ribonucleoprotein complex. J Virol 2014, 88 (13), 7602–7617. 10.1128/JVI.00256-14.

(37) Van Aalen, E. A.; Wouters, S. F. A.; Verzijl, D.; Merkx, M. Bioluminescent RAPPID Sensors for the Single-Step Detection of Soluble Axl and Multiplex Analysis of Cell Surface Cancer Biomarkers. Anal Chem 2022, 94 (17), 6548–6556. 10.1021/acs.analchem.2c00297.

(38) Nishihara, R.; Dokainish, H. M.; Kihara, Y.; Ashiba, H.; Sugita, Y.; Kurita, R. Pseudo-Luciferase Activity of the SARS-CoV-2 Spike Protein for Cypridina Luciferin. ACS Cent Sci 2024, 10 (2), 283–290. 10.1021/acscentsci.3c00887.

(39) Hunt, A. C.; Case, J. B.; Park, Y.-J.; Cao, L.; Wu, K.; Walls, A. C.; Liu, Z.; Bowen, J. E.; Yeh, H.-W.; Saini, S.; Helms, L.; Zhao, Y. T.; Hsiang, T.-Y.; Starr, T. N.; Goreshnik, I.; Kozodoy, L.; Carter, L.; Ravichandran, R.; Green, L. B.; Matochko, W. L.; Thomson, C. A.; Vögeli, B.; Krüger, A.; VanBlargan, L. A.; Chen, R. E.; Ying, B.; Bailey, A. L.; Kafai, N. M.; Boyken, S. E.; Ljubetič, A.; Edman, N.; Ueda, G.; Chow, C. M.; Johnson, M.; Addetia, A.; Navarro, M.-J.; Panpradist, N.; Gale, M.; Freedman, B. S.; Bloom, J. D.; Ruohola-Baker, H.; Whelan, S. P. J.; Stewart, L.; Diamond, M. S.; Veesler, D.; Jewett, M. C.; Baker, D. Multivalent Designed Proteins Neutralize SARS-CoV-2 Variants of Concern and Confer Protection against Infection in Mice. Sci Transl Med 2022, 14 (646). 10.1126/scitranslmed.abn1252.

(40) Gräwe, A.; Merkx, M. Bioluminescence Goes Dark: Boosting the Performance of Bioluminescent Sensor Proteins Using Complementation Inhibitors. ACS Sens 2022, 7 (12), 3800–3808. 10.1021/acssensors.2c01726.

(41) Keeble, A. H.; Turkki, P.; Stokes, S.; Anuar, I. N. A. K.; Rahikainen, R.; Hytönen, V. P.; Howarth, M. Approaching Infinite Affinity through Engineering of Peptide-Protein Interaction. Proc Natl Acad Sci U S A 2019, 116 (52), 26523–26533. 10.1073/pnas.1909653116.

(42) Tsang, T. K.; Cowling, B. J.; Fang, V. J.; Chan, K.-H.; Ip, D. K. M.; Leung, G. M.; Peiris, J. S. M.; Cauchemez, S. Influenza A Virus Shedding and Infectivity in Households. Journal of Infectious Diseases 2015, 212 (9), 1420–1428. 10.1093/infdis/jiv225.

(43) Transfiguracion, J.; Manceur, A. P.; Petiot, E.; Thompson, C. M.; Kamen, A. A. Particle Quantification of Influenza Viruses by High Performance Liquid Chromatography. Vaccine 2015, 33 (1), 78–84. 10.1016/j.vaccine.2014.11.027.

(44) Syed, A. J.; Anderson, J. C. Applications of Bioluminescence in Biotechnology and Beyond. Chem Soc Rev 2021, 50 (9), 5668–5705. 10.1039/D0CS01492C.

(45) Yang, Y.; Du, L.; Liu, C.; Wang, L.; Ma, C.; Tang, J.; Baric, R. S.; Jiang, S.; Li, F. Receptor Usage and Cell Entry of Bat Coronavirus HKU4 Provide Insight into Bat-to-Human Transmission of MERS Coronavirus. Proceedings of the National Academy of Sciences 2014, 111 (34), 12516–12521. 10.1073/pnas.1405889111.

(46) Tang, N. C.; Chilkoti, A. Combinatorial Codon Scrambling Enables Scalable Gene Synthesis and Amplification of Repetitive Proteins. Nat Mater 2016, 15 (4), 419–424. 10.1038/nmat4521.

(47) Rijal, P.; Wang, B. B.; Tan, T. K.; Schimanski, L.; Janesch, P.; Dong, T.; McCauley, J. W.; Daniels, R. S.; Townsend, A. R.; Huang, K.-Y. A. Broadly Inhibiting Antineuraminidase Monoclonal Antibodies Induced by Trivalent Influenza Vaccine and H7N9 Infection in Humans. J Virol 2020, 94 (4). 10.1128/JVI.01182-19.

(48) Sachs, L. Applied Statistics; Springer New York: New York, NY, 1984. 10.1007/978-1-4612-5246-7.

