## supporting_information for "Direct and ultrasensitive bioluminescent detection of intact respiratory viruses"

**This PDF file includes:**

Figures S1 to S12
Section S1 Supplementary Materials and Methods
Section S2 Protein Sequences

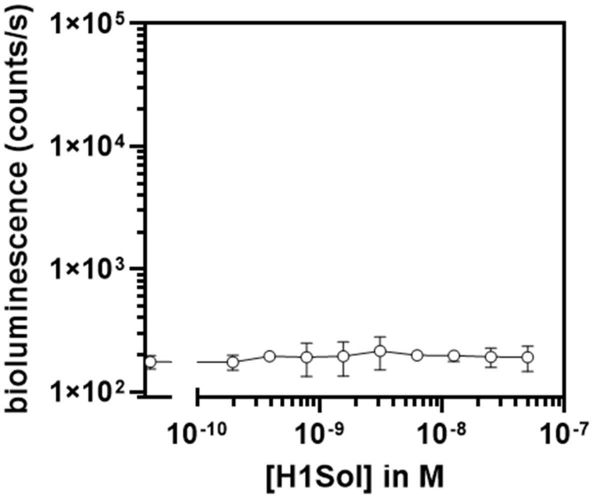


**Figure S1.**

**GLOVID without binders attached**.

LgBiT-Dog1 and SmBiT-Dog1 were mixed and used in titrations agains soluble, trimeric viral surface protein H1Sol (H1 from A/Solomon Islands/3/2006 (H1N1). Experimental conditions: 5 nM LgBiT-Dog1, 5 nM SmBiT-Dog1, final NanoGlo dilution 1:1000, 1 h incubation 22 °C. Error bars represent the standard deviation of n=3 replicates.

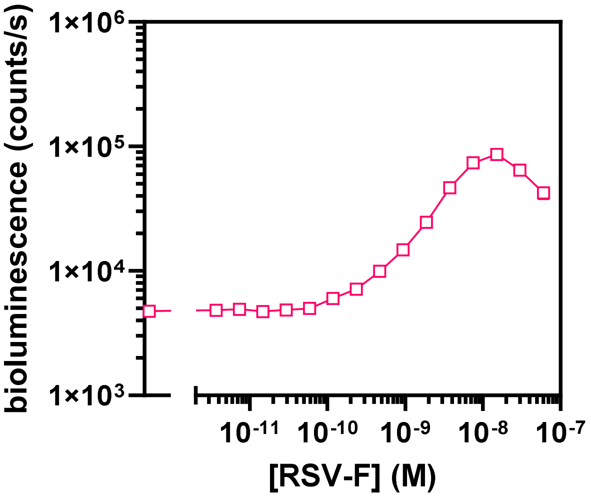


**Figure S2.**

**Trivalent GLOVID with F-VHH-4.**

GLOVID with anti-prefusion RSV-F nanobody F-VHH-4 conjugated to LgBiT-Dog3 and SmBiT-Dog3 (trivalent system). Experimental conditions: 1 nM GLOVID components, 1xPBS + 1mg/ml BSA, final NanoGlo dilution 1:2000, 1 h incubation 22 °C. The mean of n=3 replicates is shown, with error bars too small to be depicted.

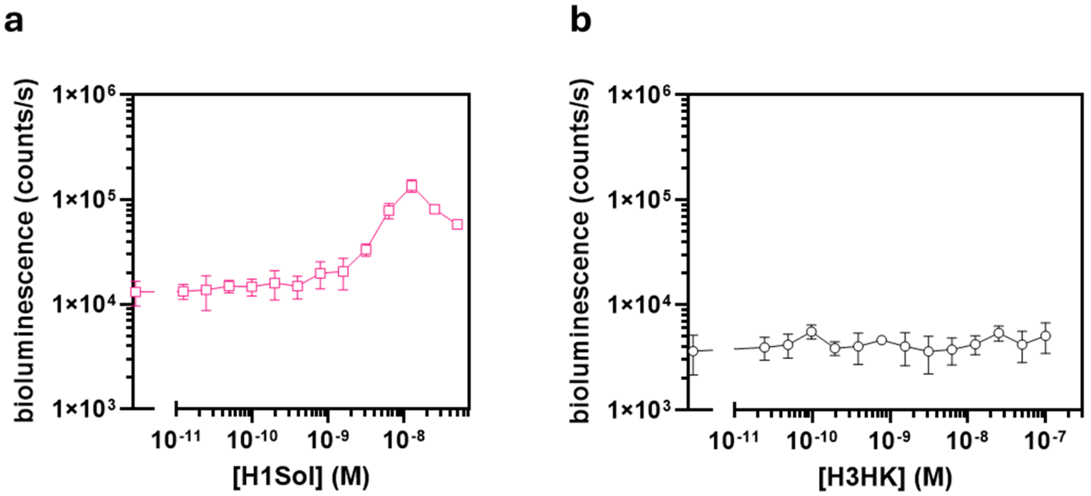


**Figure S3.**

**Trivalent GLOVID and control on H3HK with S5.**

a) Trivalent GLOVID where cyclic peptide S5 was conjugated to each LgBiT-Dog3 and SmBiT-Dog3 and used in titrations of H1Sol. b) Control GLOVID on H3HK with cyclic peptide S5 conjugated to both sensor components. Experimental conditions: 2 nM LgBiT, 6 nM SmBiT, 1xPBS, 1 h incubation at 22 °C, 1:1000 diluted NanoGlo. Error bars represent the standard deviation of n=3 replicates.

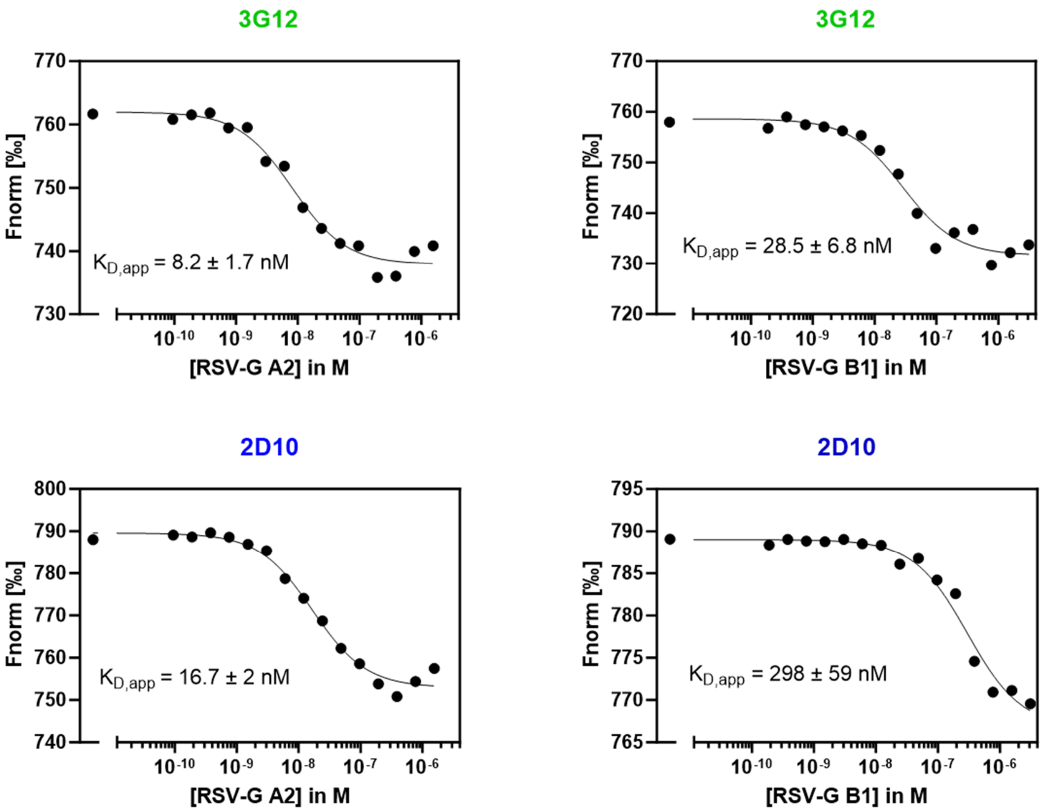


**Figure S4.**

**MST of scFv variants targeting RSV-G**

Microscale thermophoresis (MST) experiments for anti-RSV-G scFv versions of 3G12 and 2D10. A final concentration of Alexa647-labelled scFvs of 2 nM was used, with 60% excitation power and 40% MST power. MST on anti-N2 AS4C-HL-DogTag was previously described in (*1*). Shown are traces from single experiments.

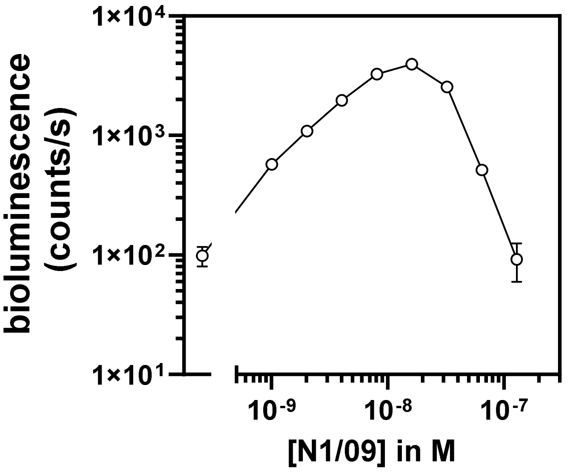


**Figure S5.**

**Monovalent GLOVID with 1GO1 scFv**

GLOVID assays with scFv binder 1GO1 conjugated to LgBiT-Dog1 and SmBiT-Dog1, targeting IAV N1/09. Experimental conditions: final GLOVID component concentration 2 nM, 1xPBS + 1 mg/ml BSA, 16 h incubation at 4 °C, final NanoGlo dilution 1:2000. Error bars represent the standard deviation of n=3 replicates.

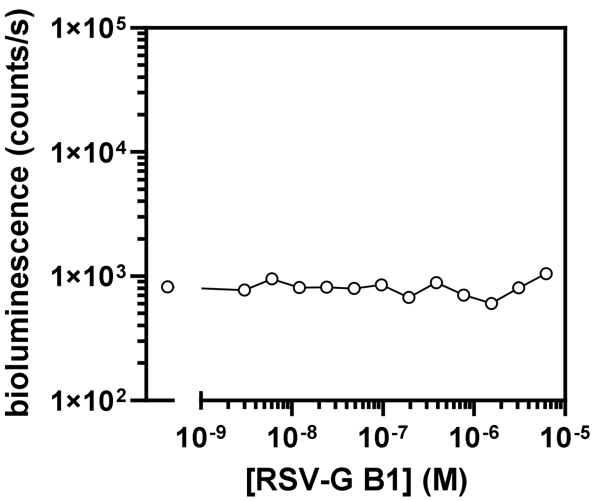


**Figure S6.**

**Controls of anti RSV-G GLOVID (1/2)**

GLOVID on RSV-G B1 assay using a combination of 3G12-LgBiT and 2D10-SmBiT. Experimental conditions: 1xPBS plus 1 mg/ml BSA, 2 nM each sensor component, 1 h incubation at 22 °C, 1:2000 diluted NanoGlo substrate. The mean of n=3 replicates is shown, with error bars too small to be depicted.

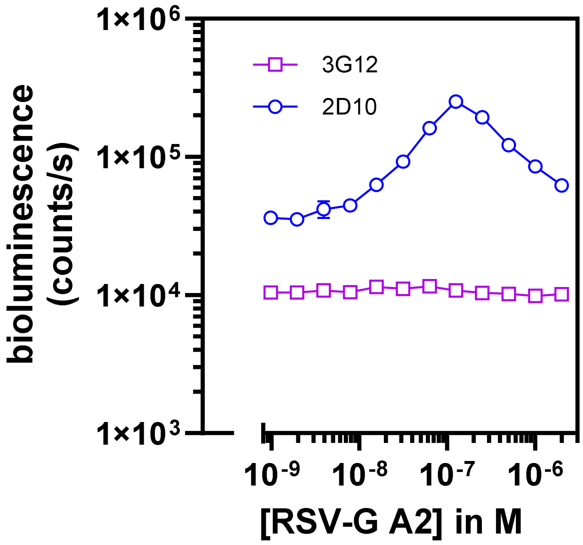


**Figure S7.**

**Controls of anti RSV-G GLOVID (2/2)**

GLOVID on RSV-G A2 using the same scFv (3G12 or 2D10) on both sensor components. Experimental conditions: 1xPBS plus 1 mg/ml BSA, 4 nM each sensor component, incubation 4 °C for 16 h to reach full binding equilibrium, 1:2000 diluted NanoGlo substrate. Error bars represent the standard deviation of n=3 replicates.

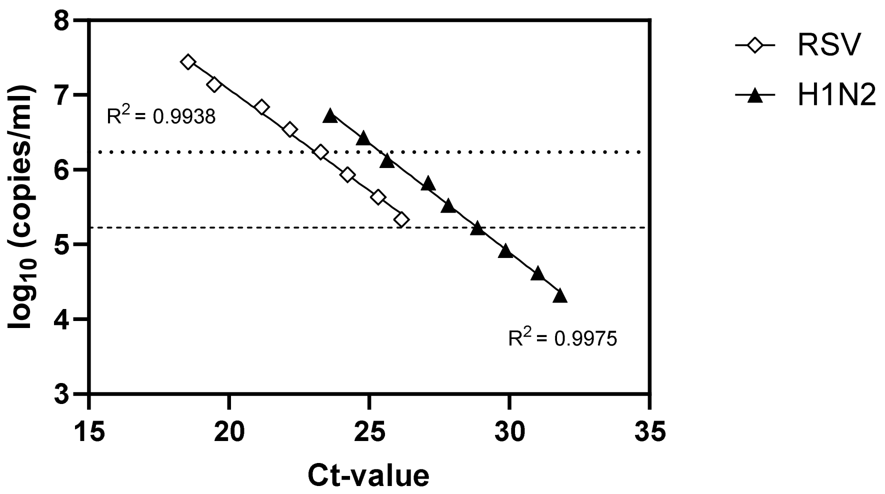


**Figure S8.**

**RT-qPCR standard curves**

Standard curves of RT-qPCR on H1N2 (GeneXpert Flu/SARS/RSV triplex plus) and RSV-A2 (Seegene RV essential). The dashed line corresponds to the LoD of the H1N2 GLOVID (Ct 28.86); the dotted line corresponds to the LoD of the RSV-A2 GLOVID (Ct 23.27). Data points represent single measurements from a dilution series of H1N2 or RSV-A2, respectively.

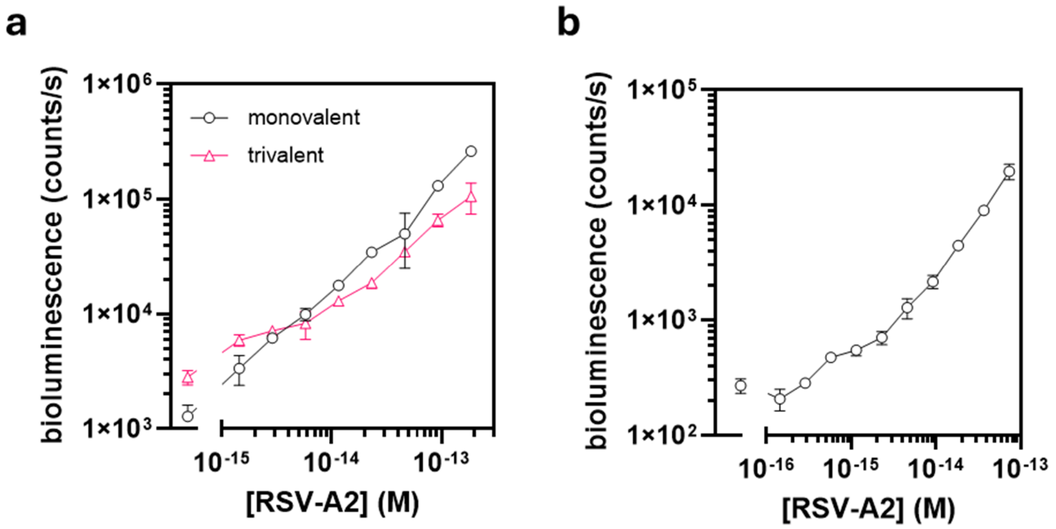


**Figure S9.**

**GLOVID assays targeting RSV-A2 via one surface protein.**

a) GLOVID assay that uses F-VHH-4 as binder, in monovalent fashion (F-VHH-4 fused to LgBiT-Dog1 and SmBiT-Dog1) and trivalent fashion (F-VHH-4 fused to LgBiT-Dog3 and SmBiT-Dog3); b) GLOVID assay that uses 3G12 (fused to LgBiT-Dog1) and 2D10 (fused to SmBiT-Dog1) as sensor parts. Experimental conditions: 4 nM final GLOVID component concentration, 1xPBS, final NanoGlo dilution 1:2000, 2 h incubation 22 °C. Error bars represent the standard deviation of n=3 replicates.

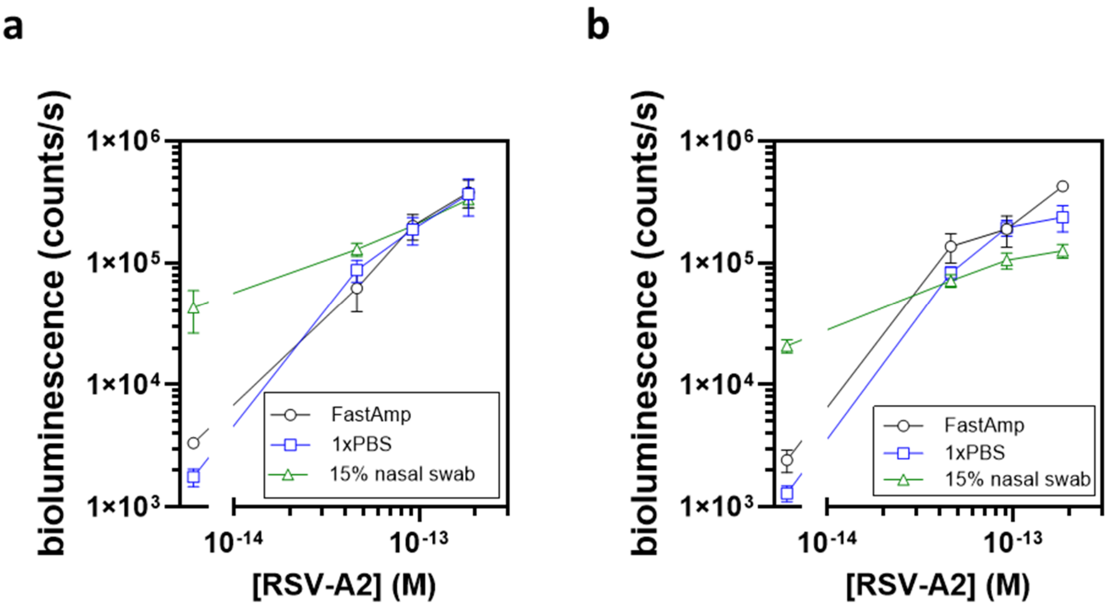


**Figure S10.**

**Spiking experiments in different matrices.**

Spiking experiment where virus (RSV-A2) was added to different matrices (FastAmp (Intact Genomics), 1xPBS, 15% diluted nasal swab) and tested with GLOVID via F-VHH-4-LgBiT / F-VHH-4-SmBiT (a) or 2D10-LgBiT / F-VHH-4-SmBiT (b). Experimental conditions: 3 nM LgBiT, 6 nM SmBiT, buffer as indicated in the legend, final NanoGlo dilution 1:1000, 1 h incubation at 22 °C. Error bars represent the standard deviation of n=3 replicates.

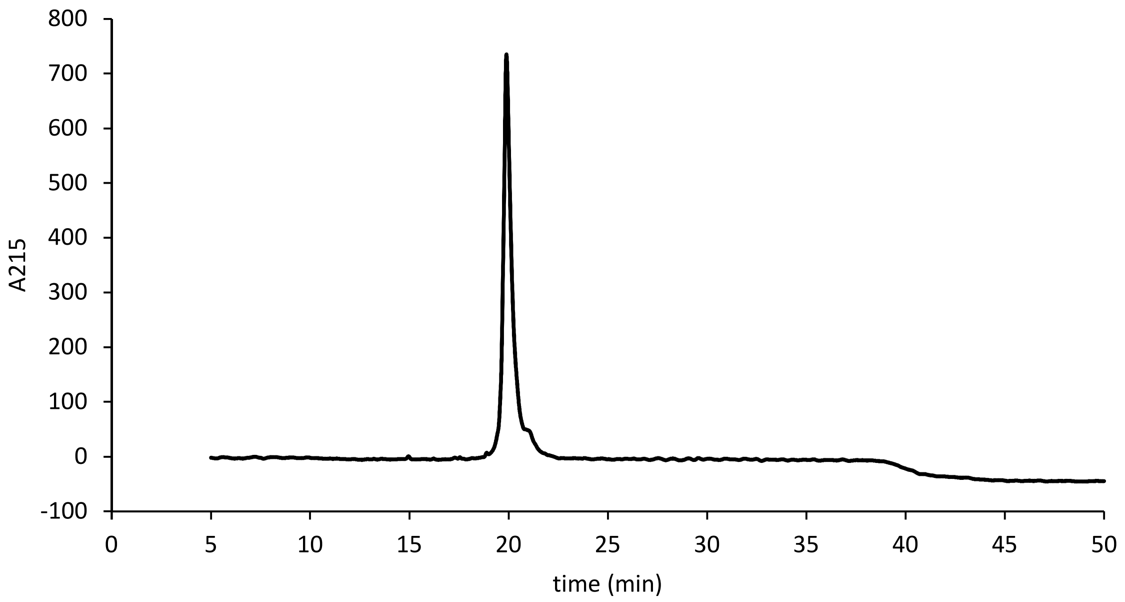


**Figure S11.**

**Analytical HPLC trace of purified final S5-dog tag product.**

The purification was performed on a C18 column, with an elution gradient 27.5-47.5% v/v acetonitrile in water with 0.1% trifluoroacetic acid.

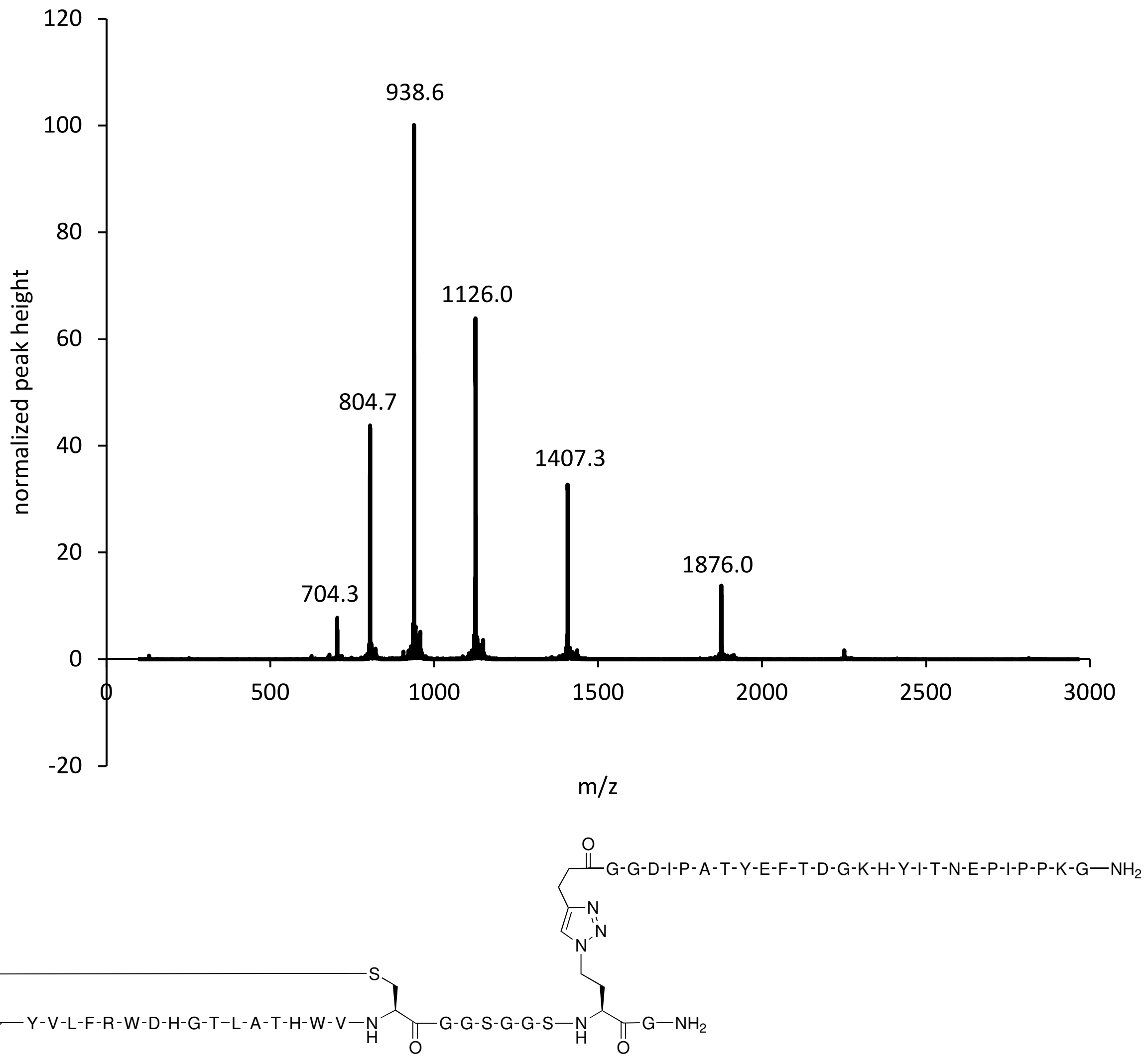


**Figure S12.**

**Mass spectrum of HPLC peak at retention time 19.9 min (Figure S13).**

Calculated for final product $C_{255}H_{364}N_{70}O_{74}S$: 5622.66, found 804.7 (calc. for $[M+7H]^{7+}$: 804.2), 938.6 (calc. for $[M+6H]^{6+}$: 938.1), 1126.0 (calc. for $[M+5H]^{5+}$: 1125.5), 1407.3 (calc. for $[M+4H]^{4+}$: 1406.7), 1876.0 (calc. for $[M+3H]^{3+}$: 1875.2).

### Section S1 – Supplementary Materials and Methods

#### Microscale Thermophoresis (MST) experiments

To estimate the $K_D$ of the constructed 2D10-HL-DogTag and 3G12-HL-DogTag scFvs for binding to RSV-G variants, purified scFvs were labeled with Alexa647 and used in MST binding experiments. For labelling, the protein was concentrated using an Amicon filter (10 MWCO) and buffer exchanged to 0.2 M sodium bicarbonate pH 8.3 using PD SpinTrap G-25 columns (Cytiva) according to the manufacturer's protocol. 100 µl of each protein at ~15 µM was mixed with 10 µl of Alexa-647 NHS Ester (Lumiprobe) freshly dissolved in DMSO (10 mg/ml) and incubated for 1 h at 22 °C with constant shaking. The reaction was purified from excess dyes by subsequently applying it to PD SpinTrap G-25 column twice. The concentration of the protein and the efficiency of labeling was calculated according to (2). Labelled scFvs were mixed with varying concentrations of target and incubated for 1 h at 22 °C in a volume of 40 µl in 1xPBS & 0.02% Tween20 (final scFv concentration 2 nM). Capillaries were loaded and MST experiment was performed at 60% LED power and 40% MST power on a Monolith NT.115 (NanoTemper Technologies).

#### ddPCR

Viral RNA from virus samples was prepared for digital-droplet PCR (ddPCR) by adding an inactivation buffer (200 mM TCEP, 2 mM EDTA, 2 U/µl murine RNase inhibitor, 20 mM Tris-HCl, pH 8.0) to the sample in 1:1 ratio, followed by incubation at 95°C for 5 min. Subsequently, the 1-Step RT-ddPCR Advanced Kit for Probes (BioRad) was used in combination with the CFX96 thermocycler and the QX200 ddPCR system, using the following oligos and probes depending on the virus targeted. Data was analysed using BioRad QX One (v1.2). The oligos target the M gene of IAV and the M gene of RSV, respectively.

**ddPCR oligo list**

| IAV targeting oligos | sequences (5' → 3') |
|---|---|
| FLUAM-7-F | CTTCTAACCGAGGTCGAAACGTA |
| FLUAM-161-R | GGTGACAAGATTGGTCTTGTCTTTA |
| FLUAM-49-P | TCAGGCCCCCTCAAAGCCGAG |
| RSV targeting oligos | |
| RSVM-F | GGCAAATATGGAAACATACGTGAA |
| RSVM-R | TCTTTTTCTAGGACATTGTATTGAACAG |
| RSVM-P | CTGTGTATGTGGAGCCTTCGTGAAG |

#### Influenza A virus and RSV sample collection from residual clinical materials

Patients were swabbed with eSwab (Copan, Italy) flocked tips, containing 1 ml of liquid modified Amies fluid. Molecular diagnostics were performed using GeneXpert SARS-CoV-2/Flu/RSV rapid test (Cepheid, Sunnyvale, CA, USA), BioFire RP2.1plus rapid test (bioMérieux, France), and Allplex RV Essential assay RT-PCR (Seegene, Seoul, South Korea) combined with the FlowGO middle ware (LabHelp Labautomation, Bladel, the Netherlands) as previously described (*3*). Standard curves for correlating the ddRT-PCR experiments to GeneExpert and Seegene Ct-values were obtained from dilution series of the A/swine/Italy/114922/2014 (H1N2) and RSV-A2 stocks.

**Section S2**
**Protein sequences**

The used tags for protein purifications are hexa-His-tag (HHHHHH) and Strep-tag II (WSHPQFEK).

LgBiT-DogCatcher (LgBiT-Dog1):
Blue: LgBiT, Red: DogCatcher
MGTSVFTLEDFVGDWEQTAAYNLDQVLEQGGVSSLLQNLAVSVTPIQRIVRSGENALKI
DIHVIIPYEGLSADQMAQIEEVFKVVYPVDDHHFKVILPYGTLVIDGVTPNMLNYFGRPY
EGIAVFDGKKITVTGTLWNGNKIIDERLITPDGSMLFRVTINSSGGGTKLGEIEFIKVDKTD
KKPLRGAVFSLQKQHPDYPDIYGAIDQNGTYQDVRTGEDGKLTFTNLSDGKYRLIENSEP
PGYKPVQNKPIVSFRIVDGEVRDVTSIVPQGGGSWSHPQFEK*

DogCatcher-SmBiT (SmBiT-Dog1):
Red: DogCatcher, yellow: SmBiT101 (*4*)
MGTKLGEIEFIKVDKTDKKPLRGAVFSLQKQHPDYPDIYGAIDQNGTYQDVRTGEDGKL
TFTNLSDGKYRLIENSEPPGYKPVQNKPIVSFRIVDGEVRDVTSIVPQGKLGGSGGSGGSG
GSGGSGGSGGSGGSGGENLYFQSGGSGGSGGSGGSGGSGGSGGSGGTGSVTGYRLFEKE
SGGSGGSWSHPQFEK*

SD36-DogTag:
Blue: SD36, green: DogTag.
MGSVQLVESGGGLVQAGGSLKLSCAASGRTYAMGWFRQAPGKEREFVAHINALGTRTY
YSDSVKGRFTISRDNAKNTEYLEMNNLKPEDTAVYYCTAQGQWRAAPVAVAAEYEFW
GQGTQVTVSGGSGGSGTGDIPATYEFTDGKHYITNEPIPPKGGSGGSWSHPQFEK*

HSB2.A-DogTag:
Blue: HSB2.A, green: DogTag.
MGSHHHHHHSSGGIVNVPNCNTTKYQQLARTAVAIYNYHEQAHLTFVENLNCKEQGNY
YYITLAATDDAGKKAIYEAKIGVVESAGWTGVEEFKLVGSGGSGGSGGSGGSGGSSGGS
GSGTGDIPATYEFTDGKHYITNEPIPPKGGSGGSWSHPQFEK*

SD38-DogTag:
Blue: SD38, green: DogTag.
MEVQLVESGGGLVQPGGSLRLSCAVSISIFDIYAMDWYRQAPGKQRDLVATSFRDGSTN
YADSVKGRFTISRDNAKNTLYLQMNSLKPEDTAVYLCHVSLYRDPLGVAGGMGVYWG
KGALVTVSSKLGGSGGSGGSGGSGGSTGDIPATYEFTDGKHYITNEPIPPKGGSGGSWSH
PQFEK*

F-VHH-4-DogTag:
Blue: F-VHH-4, green: DogTag
MGSQVQLQESGGGLVQPGGSLRLSCAASGFTLDYYYIGWFRQAPGKEREAVSCISGSSGS
TYYPDSVKGRFTISRDNAKNTVYLQMNSLKPEDTAVYYCATIRSSSWGGCVHYGMDYW
GKGTQVTVSSKLGGSGGSGGSGGSGGSTGDIPATYEFTDGKHYITNEPIPPKGGSGGSWS
HPQFEK*

LgBiT-3xDogCatcher (LgBiT-Dog3)
Blue: LgBiT, red: DogCatcher (3 times)

MGTSVFTLEDFVGDWEQTAAYNLDQVLEQGGVSSLLQNLAVSVTPIQRIVRSGENALKI
DIHVIIPYEGLSADQMAQIEEVFKVVYPVDDHHFKVILPYGTLVIDGVTPNMLNYFGRPY
EGIAVFDGKKITVTGTLWNGNKIIDERLITPDGSMLFRVTINGGSGELTGGSGGSGGSGGS
GGSGGSGEFAEAAAKEAAAKEAAAKEAAAKEAAAKEAAAKAEFGGSGGSGGSGGSGG
SGGSGGTKLGEIEFIKVDKTDKKPLRGAVFSLQKQHPDYPDIYGAIDQNGTYQDVRTGED
GKLTFTNLSDGKYRLIENSEPPGYKPVQNKPIVSFRIVDGEVRDVTSIVPQTSTPTPTPTPTP
TPTPTPTPTPTPTPTPTPTPTPTPTPTPTPTPTPASKLGEIEFIKVDKTDKKPLRGAVFSLQKQHP
DYPDIYGAIDQNGTYQDVRTGEDGKLTFTNLSDGKYRLIENSEPPGYKPVQNKPIVSFRIV
DGEVRDVTSIVPQTGPTPTPTPTPTPTPTPTPTPTPTPTPTPTPTPTPTPTPTPTPKLGEIEFIK
VDKTDKKPLRGAVFSLQKQHPDYPDIYGAIDQNGTYQDVRTGEDGKLTFTNLSDGKYRL
IENSEPPGYKPVQNKPIVSFRIVDGEVRDVTSIVPQGGGSWSHPQFEK*

3xDogCatcher-SmBiT (SmBiT-Dog3):
Red: DogCatcher (3 times), yellow: SmBiT101
MGTKLGEIEFIKVDKTDKKPLRGAVFSLQKQHPDYPDIYGAIDQNGTYQDVRTGEDGKL
TFTNLSDGKYRLIENSEPPGYKPVQNKPIVSFRIVDGEVRDVTSIVPQTSTPTPTPTPTPTPT
PTPTPTPTPTPTPTPTPTPTPTPTPTPTPTPTPTPASKLGEIEFIKVDKTDKKPLRGAVFSLQKQHPDY
PDIYGAIDQNGTYQDVRTGEDGKLTFTNLSDGKYRLIENSEPPGYKPVQNKPIVSFRIVDG
EVRDVTSIVPQTGPTPTPTPTPTPTPTPTPTPTPTPTPTPTPTPTPTPTPTPTPTPKLGEIEFIKVD
KTDKKPLRGAVFSLQKQHPDYPDIYGAIDQNGTYQDVRTGEDGKLTFTNLSDGKYRLIE
NSEPPGYKPVQNKPIVSFRIVDGEVRDVTSIVPQGGGSGELTGGSGGSGGSGGSGGSGGS
GEFAEAAAKEAAAKEAAAKEAAAKEAAAKEAAAKAEFGGSGGSGGSGGSGGSGGSGG
TVTGYRLFEKESGGSHHHHHH*

**scFvs:**
scFv sequences are listed in their secreted form, omitting the α-factor secretion signal present in pPICZalphaB.

AS4C-HL-DogTag:
The original heavy and light chain sequences of the AS4C antibody were provided by Pramila Rijal and Alan Townsend. A heavy chain-light chain (HL) configuration scFv was designed based on these sequences.
Green: DogTag
EAEAAG(*AS4C_heavy_chain*)GGGGSGGGGSGGGGS(*AS4C_light_chain*)GTSGGGSGTGDIP
ATYEFTDGKHYITNEPIPPKGGSGGSHHHHHH*

1GO1-HL-DogTag:
Underlined: 1GO1-HL, green: DogTag
EAEAAGEVQLVESGGRALRPGGSLRLSCAASGFKFDDYAMSWVRQVPGKGLEFVSGLN
WNGDITAYTDSVKGRFTVSRDNAKNSLYLHINSPKPEDTALYYCARTSSWGDYTRGPEP
KITWYFDLWGRGTLVTVSSGGGGSGGGGSGGGGSDIQLTQSPSFLSASVGDRITITCRAS
QGIDGYLAWYQQRPGKAPNLLIYAASLLQSGVPSRFSGSGYGTEFTLTISSLQPEDFATYY
CQHLDSYPLFTFGPGTKVDIKRTGTSGGGSGTGDIPATYEFTDGKHYITNEPIPPKGGSGG
SHHHHHH*

2D10-HL-DogTag:
Underlined: 2D10-HL, green: DogTag
EAEAAGQVQLVQSGPEVKKPGASVRLSCKASGYVFTNYGVSWVRQAPGQGLEWMGWS
SPYNGNTYYAQKLKARVTMTTDTSTNTAYMELRSLRSDDTAVYYCGRDMLGVVQAVA
GPFDSWGQGTLVTVSSASGGGGSGGGGSGGGGSGGGDTPMTQSPSSVSASVGDRVTISC

RASQGISNSLAWYQQKLGKAPQLLIYAASSLQSGVPSRFSGSGSGTDFTLTISSLQPEDFA
TYYCQQTNTFPFTFGPGTKVEVRRGTSGGGSGTGDIPATYEFTDGKHYITNEPIPPKGGSG
GSHHHHHH*

3G12-HL-DogTag
Underlined: 3G12-HL, green: DogTag
EAEAAGQLQLQESGPGLVKPSETLSLTCTVSGGSISSSNYYWGWIRQPPGKGLEWIASIHD
SGSIYYNPSLRSRVTISVDTSKNQFSLKLSSVTAADTAVYYCARHLVWFGELRNNWFDP
WGQGTLVTVASGGGGSGGGGSGGGGSGGGEIVMTQSPATLSVSPGERATLSCRASQSVN
SNLAWYQHKPGQAPRLLIYGASTRATGIPARFSGSGSGTDFTLTISSLQSEDFAVYYCQQY
NNWPLFGPGTKVDLKRTGTSGGGSGTGDIPATYEFTDGKHYITNEPIPPKGGSGGSHHHH
HH*

### References


1. A. Gräwe, C. M. Spruit, R. P. de Vries, M. Merkx, Bioluminescent detection of viral surface proteins using branched multivalent protein switches. *RSC Chem Biol* **5**, 148–157 (2024).
2. J. S. Nanda, J. R. Lorsch, "Labeling a Protein with Fluorophores Using NHS Ester Derivitization" in *Methods in Enzymology*, J. R. Lorsch, Ed. (2014) vol. 536, pp. 87–94.
3. J. Flipse, A. T. Tromp, D. Thijssen, N. van Xanten-Jans-Beken, R. Pauwelsen, H. J. van der Veer, J. M. Schlaghecke, C. M. A. Swanink, Optimization of the STARlet workflow for semi-automatic SARS-CoV-2 screening of swabs and deep respiratory materials using the RealAccurate Quadruplex SARS-CoV-2 PCR kit and Allplex SARS-CoV-2 PCR kit. *Microbiol Spectr* **12** (2024).
4. A. S. Dixon, M. K. Schwinn, M. P. Hall, K. Zimmerman, P. Otto, T. H. Lubben, B. L. Butler, B. F. Binkowski, T. MacHleidt, T. A. Kirkland, M. G. Wood, C. T. Eggers, L. P. Encell, K. V. Wood, NanoLuc Complementation Reporter Optimized for Accurate Measurement of Protein Interactions in Cells. *ACS Chem Biol* **11**, 400–408 (2016).
